# MeXalign – A Segmentation Mask-Based Registration Pipeline for Multimodal Single-Cell Analysis

**DOI:** 10.64898/2026.11.13.766144

**Authors:** Claude Molitor, Lyndsey Hendriks, Max Sinner, David Loibnegger, Martin Schaier, Antonia Hafner, Walter Berger, Gunda Koellensperger

## Abstract

Leveraging the power of multimodal imaging relies on accurate coregistration of yet fundamentally different modalities. Reliable cell-to-cell correspondence is especially challenging as consecutive tissue sections do not necessarily contain identical cells or cellular structures. Here we present MeXalign, a segmentation-mask-based coregistration pipeline designed to integrate multimodal images acquired sequentially from the same specimen at single-cell resolution. Single-cell segmentation masks provide a modality-independent representation of cellular morphology and spatial organization. MeXalign was tested based on the two platforms, laser ablation inductively coupled plasma time-of-flight mass spectrometry (LA-ICP-TOFMS) and immunofluorescence (IF) microscopy. As a prerequisite, each modality integrates membrane and/or nucleus labels for accurate segmentation. The added value of MeXalign was highlighted for imaging of cell models deposited on glass slides and tissue sections. The performance of coregistration was validated on cytospins, murine tumor and human skin samples. A dedicated scoring system was introduced allowing to assess the quality of image coregistration. The robustness towards under-segmentation and downstream image analysis was evaluated on LA-ICP-TOFMS with and without coregistration, investigating metal drug uptake, barcoding and cellular phenotyping. A showcase study on cancer cells exposed to nanoparticles proved the accuracy, as the nanoparticle distribution was preserved at a subcellular level when using MeXalign. While all examples show the overall improved quality of segmentation, other benefits of complementary labelings can be anticipated, especially for cellular features unique to one modality.

Graphical Abstract

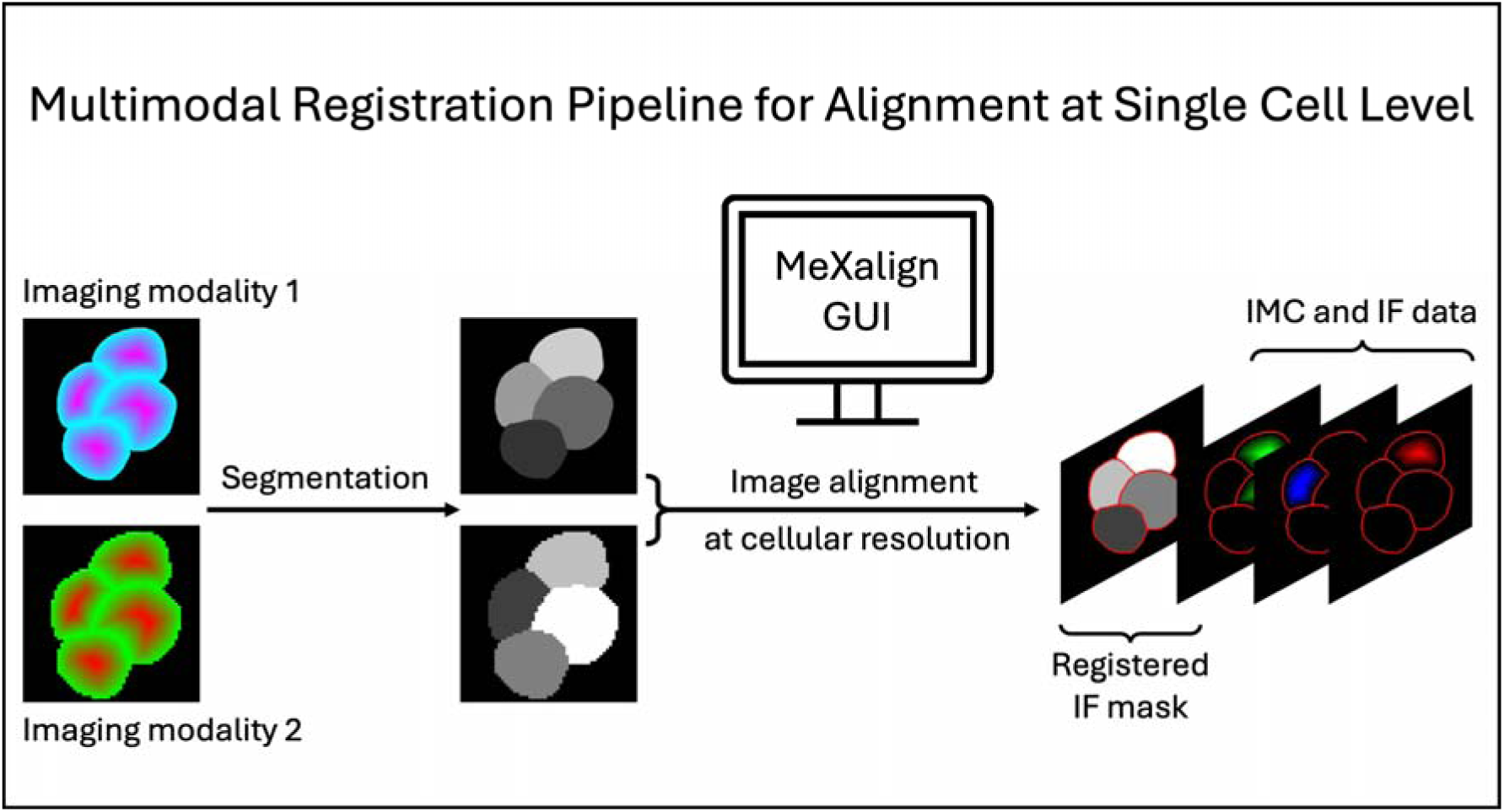

## 1. Introduction

Multimodal bioimaging is increasingly used to maximize data readout from complementary imaging modalities in the same biological specimen.^1,2^ Depending on imaging modalities used, registration pipelines operate across different scales from whole body,^3^ to tissue^4,5^ and single-cell levels.^6,7^ At each scale, a wide range of data transformations and algorithms can be used, including landmark- and silhouette-based coregistration^4,5,7^, intensity- or feature-based alignment,^6,8^ and segmentation-derived strategies.^9,10^

Traditional immunofluorescence (IF) microscopy enables fast, high-contrast acquisition of large areas at submicron spatial resolution. It benefits from well-developed marker panels, straightforward setups, and robust segmentation methods.^11^ However, standard IF is constrained by a narrow spectral range, spectral overlap and autofluorescence, limiting its multiplexing capacity. To expand readout depth, cyclic immunofluorescence (CIF)^12,13^ was introduced. Yet CIF trades off the speed and simplicity of traditional IF, requiring specialized reagents, alongside multiple consecutive cycles of staining, imaging, and bleaching. Images acquired each cycle need themselves registration. Additionally, quantification is not possible.

In contrast, mass spectrometry-based imaging methods such as imaging mass cytometry (IMC) using laser ablation inductively coupled plasma time-of-flight mass spectrometry (LA-ICP-TOFMS)^14^ enable the simultaneous detection of up to 40 metal-tagged biomarkers without spectral overlap and autofluorescence.^15^ Beyond antibodies, full-mass-range LA-ICP-TOFMS additionally captures endogenous elements, metal exposures or metal-based therapeutic agents.^16–23^ Subsequent elemental quantification enables direct comparisons of drug distributions across cells, tissue regions, and experimental groups.^16,17,23–25^ Recent hardware improvements have pushed spatial resolution from 1 µm down to 600 - 250 nm,^26–28^ providing the improved structural details required for precise segmentation and subcellular data extraction, though the lower ablation yield inherently reduces elemental sensitivity.^24,26–28^

An additional challenge in multimodal registration is that many workflows rely on adjacent or consecutive tissue sections, which do not always contain exactly the same cells or structures. While IF and IMC can be performed on consecutive slides, they can also be performed sequentially on the same tissue section, with IF acquired first followed by IMC, which is a destructive measurement. This enables complementary information from both modalities to be obtained from the same cellular structures and therefore provides a direct basis for cell-level coregistration.^6,7^

In this work, we focus on the coregistration of immunofluorescence (IF) microscopy and imaging mass cytometry (IMC), two complementary modalities that combine high-resolution imaging with highly multiplexed profiling on a cellular level. However, this pipeline can be used for every imaging modality at single cell resolution. For IF and LA-ICP-TOFMS, both can detect the same molecular targets, including DNA via intercalators and proteins via antibodies, differing only in the labels used for detection: fluorophores in IF and metal isotopes in IMC.^29^ These shared targets provide the same features for coregistration as demonstrated by Baars et al. in the MATISSE workflow.^7^ They align both modalities based on DNA-intercalator signals (DAPI from IF and Ir-DNA intercalator from IMC) using Fiji’s landmark correspondence tool with a moving-least-squares similarity transformation. In a separate approach, Kim et al likewise exploited the DNA channels for coregistration positioning multiple individual IMC measurements onto a whole-slide IF image by cross-correlation prior to stitching.^6^

These studies showed that IF and IMC are highly compatible, demonstrating that downstream analysis significantly benefits from raw-signal alignment. Working directly on the raw data avoids the information loss that comes with reducing each image to an intermediate representation (e.g. masks or sparse feature descriptors) before registration. The main difficulty, however, is that two modalities differ in signal distribution, contrast, dynamic range, background, staining variability and their spatial continuity. These differences can be further compromised by acquisition-related artifacts such as blur, saturation, and local distortions, compromising both the registration accuracy and the optimization procedures used to achieve it.

To overcome these limitations, single cell segmentation prior to registration provides an effective solution especially with recent advancements in machine learning-based segmentation^30,31^. By aligning single-cell masks rather than raw intensities, both modalities are transformed into greyscale masks ultimately eliminating signal heterogeneity and acquisition-specific differences. These masks provide a compact representation of cellular geometry and spatial organization while substantially reducing file sizes and computational load. This also allows for using full datasets compared to metric sampling methods. Coregistration then is solely dependent on segmentation performance, although poor data quality can hinder segmentation and make cell-level alignment impossible. Prior studies have successfully applied it across various scales, ranging from broad tissue features and cellular boundaries,^10^ or segmentation based strategies where segmented cells represent point sets for matching.^9^

Building on these strategies, we register IF and LA-ICP-TOFMS images directly using the label masks produced by single-cell segmentation.^30,31^ This way, cell-to-cell correspondence no longer relies on modality specific signals as proxy for convergence but directly uses defined cell masks. We adopt membrane-DNA-based segmentation as the basis for coregistration since it represents the full cell dimension. Alternatively, registration can be performed from DNA-based segmentation mask while extraction is still performed through the membrane mask as the residual deformations are spatially smooth and of low magnitude. Distortions are more relevant for large areas. Furthermore, while mask-based registration inherently depends on initial segmentation quality, we show that our workflow remains highly robust even in the presence of substantial under-segmentation. This can be achieved by transforming the label masks per stage into representations which work best for the corresponding stages. For coarse alignment, masks are binarized and each cell can be eroded independently. This creates a cellular fingerprint in featureless and background-free tissue and thereby eases FFT based translation and rotation estimation. For rigid, affine and nonrigid refinement, masks are converted into per-object-normalized gradient fields, which give a smooth cell aware landscape which optimizes convergence during intensity-driven optimization.

Subsequently, we developed MeXalign, an automated geometry-based workflow for registering segmentation masks across multimodal datasets, demonstrated here on IF to LA-ICP-TOFMS/IMC data at a single-cell resolution. Unlike conventional approaches that register measurements acquired from adjacent or consecutive tissue sections, MeXalign is designed for multimodal measurements obtained from the same physical specimen. MeXalign features a graphical user interface and combines a three-stage pipeline: (1) Fast Fourier Transform (FFT)-based coarse translation and rotation estimation, (2) rigid and affine refinement and (3) non-rigid cubic B-spline refinement for residual local deformations. The mask erosion implemented here to create a cellular fingerprint enables more reliable and accurate FFT estimation, and the single-cell gradient field-based masks developed removes modality specific differences while providing a smooth, cell-aware landscape for efficient single cell registration. All calculated transformation matrices can subsequently be applied to fluorescence channels or independent label masks.

The capabilities/performance of MeXalign was demonstrated by using it to coregister different samples, namely cytospun single cells, murine tumor tissue, and human skin tissue sections analyzed by LA-ICP-TOFMS and IF. Beyond evaluating single cell mask overlaps, we assess whether downstream analysis remains identical, mis-segmentation affects coregistration performance and if subcellular features can be transferred while preserving spatial distribution. MeXalign is thus not just a global tissue registration pipeline but a quantitatively validated framework for multimodal single-cell analysis. The pipeline is available at the KoellenspergerLab/MeXalign Github.

## 2. Experimental section

### 2.1 Sample Preparation and Measurements

#### Chemicals and Materials

Tris-buffered saline (BioUltra) and bovine serum albumin (BSA) were obtained from Sigma-Aldrich (Steinheim, Germany). Tris/EDTA-based Dako Target Retrieval Solution (pH 9) was purchased from Agilent Technologies (Waldbronn, Germany). Ultrapure water used throughout the experiments was generated in-house using a Purelab Ultra MK 2 system (ELGA, High Wycombe, U.K.). Human BD Fc Block^TM^ was supplied from BD Biosciences (San Jose, CA, USA). SuperBlock^TM^ blocking buffer in TBS and Tween 20 were obtained from Thermo Fisher Scientific (Waltham, MA, USA).

Anti-Wheat germ Agglutinin (anti-WGA) (Szabo Scandic, Vienna, Austria) and FITC-conjugated WGA (5 mg/mL) (GeneTex, Irvine, CA, USA) build the foundation for the dual label. Anti-WGA was tagged with the following lanthanides ^143^Nd, ^144^Nd, ^145^Nd, ^147^Sm, ^149^Sm, ^151^Eu, ^152^Sm, ^154^Sm, ^155^Gd, and ^165^Ho using Maxpar® X8 Antibody Labeling Kits (Standard BioTools). Concentration of metal-labeled anti-WGA were +-2 mg/ml. For DNA staining, Cell-ID^TM^ Ir-intercalator (125 µM) (Standard BioTools, San Francisco, CA, USA) and DAPI (1 µg/mL) (GeneTex, Irvine, CA, USA) were used. All other metal-tagged antibodies were acquired from Standard BioTools (San Francisco, CA, USA).

#### HCT116 cell cultivation and treatments

HCT116 wild-type (WT) human colorectal cancer cells were obtained from the American Type Culture Collection (ATCC). The corresponding oxaliplatin-resistant subline (HCT116 OxR) was established in-house by progressively increasing the oxaliplatin concentration during cultivation of HCT116 WT cells. To maintain the resistant phenotype, HCT116 OxR cells received a 10 µM oxaliplatin pulse every two weeks. Both cell lines were maintained at 37 °C and 5% CO_2_ under humidified conditions in McCoy’s 5A medium containing 10% heat-inactivated fetal calf serum (FCS; BioWest) and 2 mM L-glutamine (VWR International).

For drug-treatment experiments, cells were transferred either to T25 flasks (Greiner Bio-One) at 2 × 10^6^ cells in 5 mL medium or to T75 flasks at 5 × 10^6^ cells in 10 mL medium. Cells were allowed to adhere overnight and were subsequently exposed for 16 h to the respective treatment conditions summarized in Table S8. BOLD-100 was provided by the Institute of Inorganic Chemistry, University of Vienna, while oxaliplatin (OxPt; LC Labs; #O-7111) was obtained from LC Laboratories.

After exposure to BOLD-100, oxaliplatin, or the corresponding control conditions, cells were harvested by Trypsin/EDTA treatment (Sigma-Aldrich). Cell suspensions were centrifuged for 5 min at 1400 rpm, the supernatant was removed, and the resulting pellets were resuspended in 120 µL TBS.

For nanoparticle (10.095 mg/mL Stock; 0.49 µm Sphero^TM^ fluorescent particles Sky blue, Spherotech, Inc., Lake Forest, IL, USA) experiments, 6 × 10^5^ cells were seeded per well in 6-well plates using 2 mL medium. Cells were allowed to adhere overnight and were subsequently exposed for 24 h at 50 ng/mL. Subsequently, cells were washed with medium to remove excess nanoparticles, detached using Trypsin/EDTA and centrifuged at 1400 rpm for 5 min. The supernatant was discarded and pellet was resuspended in 120 µL TBS.

#### Optimized barcoding procedure

Barcoding was performed as described in previous work^22^ using identical reagent, concentration, incubation and staining times. In brief, approx. 1.5 x 10^6^ cells per condition were labelled with a preincubated dual label consisting of FITC-WGA and metal-labeled anti-WGA. Labels for the respective experiments are listed in Table S8 for drug treated samples and with ^155^Gd-anti-WGA for NPs treated cells. Subsequently, they were washed and pooled where stated, deposited onto Superfrost® microscope slides (Thermo Scientific) by using a cytospin cytocentrifuge (Shandon Cytospin 4, Thermo Scientific, UK), fixed with 4 % PFA and stored at - 20°C.

#### Immunostaining of Cytospins

The cytospin samples were incubated for 5 minutes at RT with a DAPI staining solution in combination with the Ir-based DNA intercalator (125 µM; Diluted 1:100 in DAPI staining solution). The sample was rinsed with ultrapure water and subsequently examined by IF and analyzed using LA-ICP-TOFMS.

#### Preparation of Human Skin tissue

The *ex vivo* skin model, NativeSkin®, from Genoskin (Toulouse, France) was used. Biopsies were obtained from surplus adult skin collected post-surgery, with all donors providing informed consent prior to donation. The use of this model was approved by the appropriate Ethics Committee in accordance with relevant guidelines and regulations. Upon arrival, the supplied culture medium was added under sterile conditions and was then incubated at 37°C with 5% CO_2_ and 95% humidity for one hour, following the manufactureŕs instructions. After treatment, collected samples were formalin-fixed, embedded in paraffin (FFPE) and 5 µm sections were prepared.

#### Preparation of murine tumor sample

The animal experiment for the tumor sample was carried out at the Institute of Cancer Research, Medical University of Vienna and was performed in accordance with the regulations of the Ethics Committee for the Care and Use of Laboratory Animals at the Medical University of Vienna (application number BMWF-66.009/0140-II/3b/2011). A female Balb/c mouse was bred in-house at the Institute of cancer Research of Vienna in February 2021. In brief, 5 x 10^5^ CT26 cells, in 50 µL serum-free RPMI-medium, were injected subcutaneously into the right flank one week prior to the first treatment (Oxaliplatin at 9 mg/kg). The first treatment was administered at an age of ∼9 weeks and repeated once after three days. Solutions were prepared freshly on the treatment days. One day after the final administration, the animal was euthanized. Tumors were collected and processed into FFPE blocks using standard procedures.

#### Immunostaining of thin tissue sections

Tissues were deparaffinized in fresh Xylene for 20 minutes and rehydrated in a graded ethanol series (from 100 % to 70 % ethanol). After washing with ultrapure water, heat mediated antigen retrieval was performed at 96°C for 30 minutes at a pH of 9 with a Tris-EDTA buffer. Samples were cooled, washed with ultrapure water and permeabilized with TBS/0.05 % Tween. Nonspecific antibody binding was reduced by incubation of the sample with SuperBlock buffer for 30 min and Human BD FC Block for 10 minutes at RT.

Membrane staining using the dual label was performed differently for skin and tissue samples. For the skin sample, FITC-WGA and ^143^Nd-anti-WGA were preincubated for 2 hours. The mixture was then applied to the tissue samples for 30 minutes. In contrast, the tumor sample was incubated with FITC-WGA alone for 30 minutes while the ^164^Dy-anti-WGA was included in the subsequent antibody cocktail.

Staining with metal-labeled antibodies was performed over night at 4 °C in a hydration chamber. The Antibody cocktails are listed within Table S1. DNA staining was performed for 5 minutes at RT with a DAPI staining solution in combination with the Ir-based DNA intercalator (125 µM; Diluted 1:100 in DAPI staining solution). Finally, the samples were washed three times with ultrapure water and air dried.

#### Fluorescence Microscopy Analysis

All samples were examined by immunofluorescence microscopy using an Axioscope 7 microscope equipped with a Colibri 7 light source (Zeiss, Oberkochen, Germany) and a 20× and a 50× objective. Images aquired using the 20× objective result in a pixel size of 0,2738 µm and 0,1010 µm for the 50× objective. IF signals were detected using the following Wavelength/Bandwidth: 469/38 nm for FITC (FITC-WGA) and 385/30 nm for DAPI.

#### LA-ICP-TOFMS Analysis

LA-ICP-TOFMS measurements were performed using an Iridia 193 nm laser ablation system (Teledyne Photon Machines, Bozeman, MT, USA) equipped with the aerosol rapid introduction system (ARIS), coupled to an ICP-TOFMS (icpTOF 2R, TOFWERK, Thun, Switzerland). Instrument performance was optimized on a daily basis using NIST SRM 612, with optimization targeting high sensitivity, minimal elemental fractionation (^238^U/^232^Th) and low oxide formation (<2.5% for ^232^Th^16^O+/^232^Th+). A detailed overview of all operating parameters is provided in Table S2. Ablation was performed with circular spot sizes of 2 - 3 μm, using an inter-spot spacing of 0.5 - 1 μm. A fixed dosage of 2 - 4 was applied, resulting in 2x to 4x overlap in both x and y directions to ensure complete material removal. Energy densities between 0.4 and 1.4 J/cm^2^ ensured full ablation of the tissues without ablating glass.

#### Data Acquisition

LA-ICP-TOFMS measurements were recorded using TofPilot software (version 2.16.2; TOFWERK AG), using the laser ablation module. Raw data were saved in HDF5 format whereas data reconstruction and handling was performed in MeXpose^23^ and MeXpose 2.0^32^ (Koellensperger Lab). Image processing was done in Fiji (v2.16.0/1.54p).^33^ Segmentation was performed using Cellpose-SAM.^31^

Coregistration was performed with MeXalign. For single-cell segmentation, nuclear signals Ir-DNA-intercalator (LA-ICP-TOFMS) and DAPI (IF) were used in combination with different membrane markers listed in Table 1. Deviations from Table 1 are noted at the respective results; for the cytospin sample coregistration was also performed once based on segmentation masks derived from DNA and ^23^Na respectively (LA-ICP-TOFMS) and DAPI (IF).

**Table 1:**
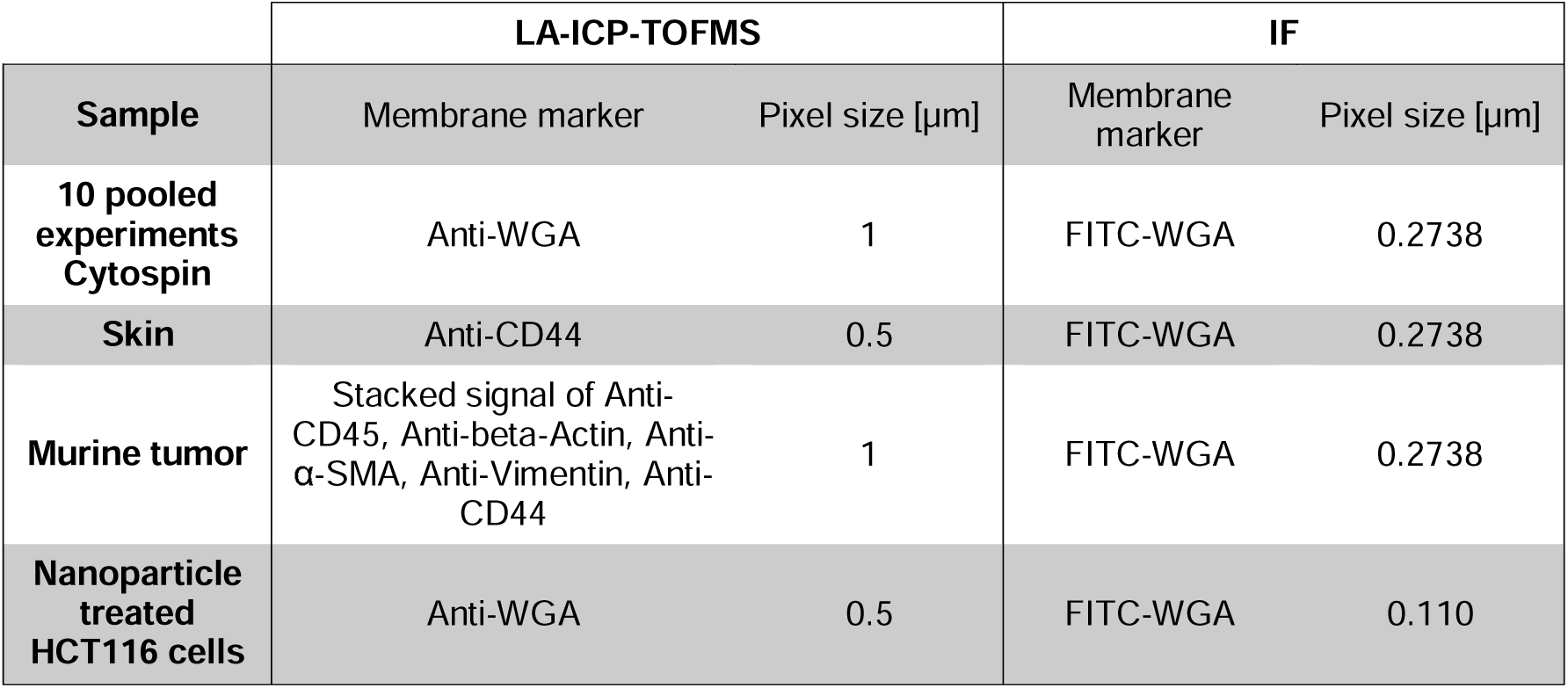
Summary of which membrane signals were used in combination with the respective DNA signals (Ir-DNA intercalator for LA-ICP-TOFMS and DAPI for IF) as well as the pixel size of the respective measurements.

### 2.2 Coregistration pipeline

#### 2.2.1 Code Basis

Image registration, image processing^34^, and supporting numerical computations were implemented in Python using SimpleITK^35^ (a simplified interface to the Insight Toolkit, ITK^36^) NumPy,^37^ SciPy,^38^ and Pillow.^39^ The graphical user interface and alignment visualizations were built with pySide^40^ and Matplotlib.^41^ Exact package versions are specified in the accompanying Conda environment file (mexalign.yml) to allow full reproduction of the software environment.

#### 2.2.2 Data preparation

The coregistration pipeline aligns IMC and IF images on a single-cell basis using segmentation-derived masks from both modalities as input (png or tiff). These are greyscale segmentation masks where each object represents a cell. While demonstrated here using IMC and IF data, the pipeline is applicable to any imaging modalities from which suitable segmentation masks can be generated. All samples within this work were segmented using Cellpose,^11,31^ though any method yielding greyscale label masks can be used.

When using label masks, background pixels are set to zero and no longer contribute to the coregistration process or its convergence. Hence, this removes modality-specific artifacts, such as out-of-focus fluorescence blur or oversaturated areas in IF, or channel smearing in IMC which could skew coregistration. Object-based masks also streamline downstream processing, enabling efficient mask erosion, binarization, vectorization, and feature extraction.

Within this workflow, the IMC images serve as fixed reference, while the IF images are the moving target. This orientation ensures that quantitative metal signals in e.g. drug uptake or environmental exposure studies remain unaltered. Therefore, IMC images dońt undergo spatial resampling or scaling during registration.

The pixel size of both modalities is fundamental for the algorithm to know the dimension differences between the modalities. Slight scaling differences can be overcome by the affine registration in stage 2. In this study, IF images have a pixel size of 0.2738 µm/pixel whereas IMC images range from 0.5 to 1.0 µm/pixel.

Additionally, a second mask can be selected to be transformed and is based on nearest-neighbor interpolation to preserve label mask integrity. All other additional IF images such as DAPI-signal will be transformed using bilinear interpolation.

#### 2.2.3 Stage 1 Fast Fourier Transformation (FFT) based coregistration

The purpose of stage 1 is to identify the approximate spatial position of the IMC image relative to the IF image. An example demonstrating FFT-based alignment^42,43^ of a small ablated region within a full cytospin sample is provided in Figure S9. The FFT alignment accounts for both coarse sample morphology and individual cellular structures. This resolution factor, the ratio between the pixel size from both modalities serves as rough scaling alignment between the two imaging modalities.

Equation 1: Ratio between the pixel size of the IMC- and IF image.

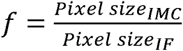

Input greyscale masks are first binarized, as identical cells carry different labels across the two modalities (Figure S9). The spatial region yielding the highest FFT cross-correlation peak is selected to determine translation. To further account for rotational misalignments, the IF image is evaluated over a user-defined angular range and step size (default ±3° in 0.25° increments), retaining the angle with the highest FFT cross-correlation peak yield. The estimated matching region is then cropped from the greyscale IF label mask according to the bounding box of the fixed IMC image, plus a safety margin (default: 100 pixels of the IF image). This padded area accommodates for minor misalignments and modality-specific distortions for refinement in later stages, while reducing data volume to accelerate subsequent computations.

In highly dense samples without extracellular background, simple binarization can cause cellular information to be lost, leaving only rough morphologies. Depending on the sample, this lack of background can cause FFT-based registration to fail. To resolve this, mask erosion can be applied to artificially generate the necessary background. By eroding the outer pixels of each object/cell before binarization of the label masks creates gaps between cells. This way, a spatial fingerprint (cell positions) of the sample is generated (Figure S8) which then enables and facilitates registration. To account for resolution differences between modalities during erosion, the erosion for the IF mask is scaled proportionally to the resolution factor (Equation 1) and is always rounded up to the next whole number of pixels. For an IMC pixel size of 1.0 µm and an IF pixel size of 0.2738 µm, eroding 1 pixel from every object in the fixed IMC mask corresponds to eroding 4 pixels (*f* =3.65 → 4) from every object in the IF mask. This ensures that the same relative distance is eroded prior to downsampling the moving mask.

#### 2.2.4 Stage 2 Rigid/Affine Refinement

The second stage establishes a global alignment across the sample. First, the cropped and padded greyscale mask is transformed to per-cell normalized gradient fields (chessboard design).^44,45^ By default, a center-based gradient is used, where each segmented object is encoded by a normalized internal distance map that peaks at the cell center and decreases gradually towards the boundary. Alternatively, an edge-based gradient can be selected; this places peak intensities at object borders and minimum values at the center, emphasizing the membrane matching. Both methods provide a well-defined cellular landscape to improve convergence even when initial overlap is limited or distorted.

Alignment is based on minimizing the mean squared (MSQ) metric, i.e. the mean of the squared intensity differences between the fixed and moving gradient fields using regular step gradient descent optimization.^36^ A rigid Euler2D transformation (translation and rotation only) is performed, followed by an affine transformation (additionally incorporating scaling and shearing) to compensate for minor magnification discrepancies and modality-dependent shear distortions.^36^ To iteratively resolve distortions from large to fine scales, affine registration is performed across a multi-resolution image pyramid.^36,46,47^ Tunable parameters for this stage include the number of iterations, learning rate and metric sampling percentage for the rigid and affine stage, the shrink factor for affine stage (see Table S3 for further details and optimization strategies). The shrink factor for the rigid stage is hardcoded as [8,4,2] pyramid. Each pyramid level stops early once the optimizer step falls below a minimum, the gradient magnitude drops below tolerance, or, for the final affine level, the MSQ metric plateaus over a set iteration window. Otherwise, it runs to the maximum iteration count. The gradient fields generation can be selected within the input tab.

#### 2.2.5 Stage 3 Nonrigid B-Spline refinement

To compensate for remaining local deformations, a cubic B-Spline free-form-deformation (FFD)^48^ field is applied to the same single-cell gradient field masks from Stage 2, initialized with the transformation parameters from Stage 2. The image domain is subdivided by a control point mesh. Displacing a control point mainly affects its own neighborhood. This results in local corrections without discontinuities. To remain independent of pixel size, the control point mesh is derived from the physical dimensions of the fixed image using a target control-point spacing (default: 150 µm for MSQ and 200 for Mattes mutual information, MMI). The physical size of each image axis is calculated by multiplying the image dimensions (in pixels) by the fixed-image pixel size. Dividing this value by the target spacing and rounding up gives the number of mesh intervals along that axis. For example, a fixed image of 3208 × 950 pixels at 0.5 µm pixel size has physical dimensions of 1604 × 475 µm, and with a spacing of 150 µm, this results in an 11 × 4 mesh, which corresponds to 14 × 7 control points for a cubic B-Spline. The deformation field is optimized across a multi-resolution image pyramid^36,46,47^ using either the mean square (MSQ) difference or Mattes mutual information (MMI)^36,46,47,49^ metric as the similarity measure alongside progressive control point refinement. Optimization is driven by a limited memory quasi-Newton optimizer (L-BFGS)^50^ whose integrated line search automatically selects the step length and iteration. Tunable parameters include the maximum number of iterations, the similarity metric (MSQ or MMI), the target control-point spacing (µm), the shrink and mesh scale factors, the early-stop window and tolerance for when convergence plateaus (Table S3). Registration quality is quantified by calculating the Dice similarity coefficient^51,52^ and Intersection over Union (IoU)^51,53^ between the fixed and registered moving mask.^54^ Folding is assessed via the Jacobian determinant of the complete applied transformation (Affine and B-spline).^55^ It is evaluated within the cell foreground of the fixed mask, excluding a 2-pixel border affected by finite-difference edge effects. Because the affine part scales the image, the values are centered on its determinant rather than on 1. FFT^43^ localization and all deformations from rigid, affine^36^ and B-Spline^48^ transform are then applied to the selected channels.

## 3. Results and Discussion

### 3.1 Coregistration validation metrics

Two common validation metrics used for comparing masks, tissue or segmentation masks are the Dice coefficient and Intersection over Union (IoU, also named Jaccard Index). Both metrics range from 0, indicating no overlap, to 1, indicating perfect overlap. For the same pair of masks, the Dice coefficient is always equal to or greater than the corresponding IoU value. The Dice coefficient is calculated as twice the overlapping area of two masks divided by the sum of their individual areas. The IoU is calculated as the overlapping area of two masks (intersection) divided by the combined area (Union) (see Figure S1). These however come with caveats as one of the major purposes of a coregistration is the improved segmentation.^6,7^ Improved segmentation means being able to segment cells which previously could not be segmented or the reduction of multiplets, so two or more cells which could not be differentiated and were segmented as one cell even though consisting of multiple. A detailed version of the modified Dice and IoU score to make segmentation comparable (Figure S6) and other alternatives are described in the supporting information and illustrated in Figure S2 - Figure S5. The modified Dice coefficient and IoU implemented here is the complete/modified score.

### 3.2 Applied study cases

To demonstrate the applicability of MeXalign and evaluate its performance, we used different biological samples including cytospun single cells and complex tissue sections. Cytospun single cells provided undense single cells and were first used as a simplified and controlled model system to systematically evaluate the requirements for successful coregistration. We tested whether registration requires the same membrane label to be present in both modalities, as in the dual WGA/anti-WGA labelling strategy, or whether it is sufficient to label the membrane using different markers, such as WGA in IF and CD44 in LA-ICP-TOFMS. We further investigated whether endogenous elemental signals can provide sufficient cellular information for coregistration without additional membrane staining. These experiments establish which types of signals and segmentation masks can be used for reliable cell-level registration. We then applied MeXalign to tissue and cytospin samples to demonstrate the practical benefits of transferring high-resolution cellular segmentation to quantitative multimodal imaging, including subcellular localization of fluorescent nanoparticles (NPs).

#### 3.2.1 Validation of the coregistration using the same dual-label signal

To establish and validate the coregistration pipeline, we first used cytospun single cells as a simplified model system. Ten different experiments were conducted using a HCT116 wild type (WT) and an Oxaliplatin resistant (OxR) cell lines. Cells were exposed to different drug treatments, and concentrations (16h with 1 or 5 µM Oxaliplatin; 5 or 100µM for BOLD-100) together with the corresponding untreated controls. Each experiment was tagged with a unique identifier, a barcode which enables pooling of all experiments and the simultaneous measurement of all experiments while still being able to retrace all cells to their respective experiment.^22^ This setup enables comparing debarcoding (unsupervised Leiden clustering by PhenoGraph) and single cell data extracted from the ICP- and the coregistered IF mask. The barcoding scheme used was published in a previous work using a WGA/Anti-WGA dual label.^22^ The benefit of this method lies within the dual label, WGA being labeled with a fluorophore and Anti-WGA labeled with lanthanoids. This enables to tag cellular membranes within IF as well as IMC and builds the foundation for segmentation and ultimately the coregistration pipeline.^26^ While the WGA / Anti-WGA dual labelling approach is quite advantageous, segmentation can be performed on other membrane markers too and will be discussed in the following sections (e.g. Figure 4 and Figure S12) or even DNA.

For a proper segmentation, the IF image within Figure S9 was cropped. The full coregistration is illustrated within Figure S10 and coregistration parameters within Table S6. To compare the coregistration, single cell data was extracted once via the normal ICP segmentation mask, and once via the coregistered IF mask (Figure S10). Cells were debarcoded (traced back to the respective experiment) using the barcode signals as input for unsupervised Leiden clustering by PhenoGraph implemented within MeXpose 2.0.^32^ Figure 2 shows the UMAP for ICP mask extracted single cell data as well as for coregistered IF mask extracted single cell data. Cells showing no signal and not being assigned to any of the ten clusters were removed as described within previous work.^22^ Both datasets show accordingly 10 distinct clusters representing the 10 experiments also in accordance with the UMAP (Figure 2). To illustrate the robustness of the data extraction on a single cell basis, the ^102^Ru, ^196^Pt and ^154^Sm (barcode signal for WT_OxP5) marker signals were compared within the respective WT sample with the highest drug concentrations. Abbreviations for the respective samples are listed in Table S8.

**Figure 1:**
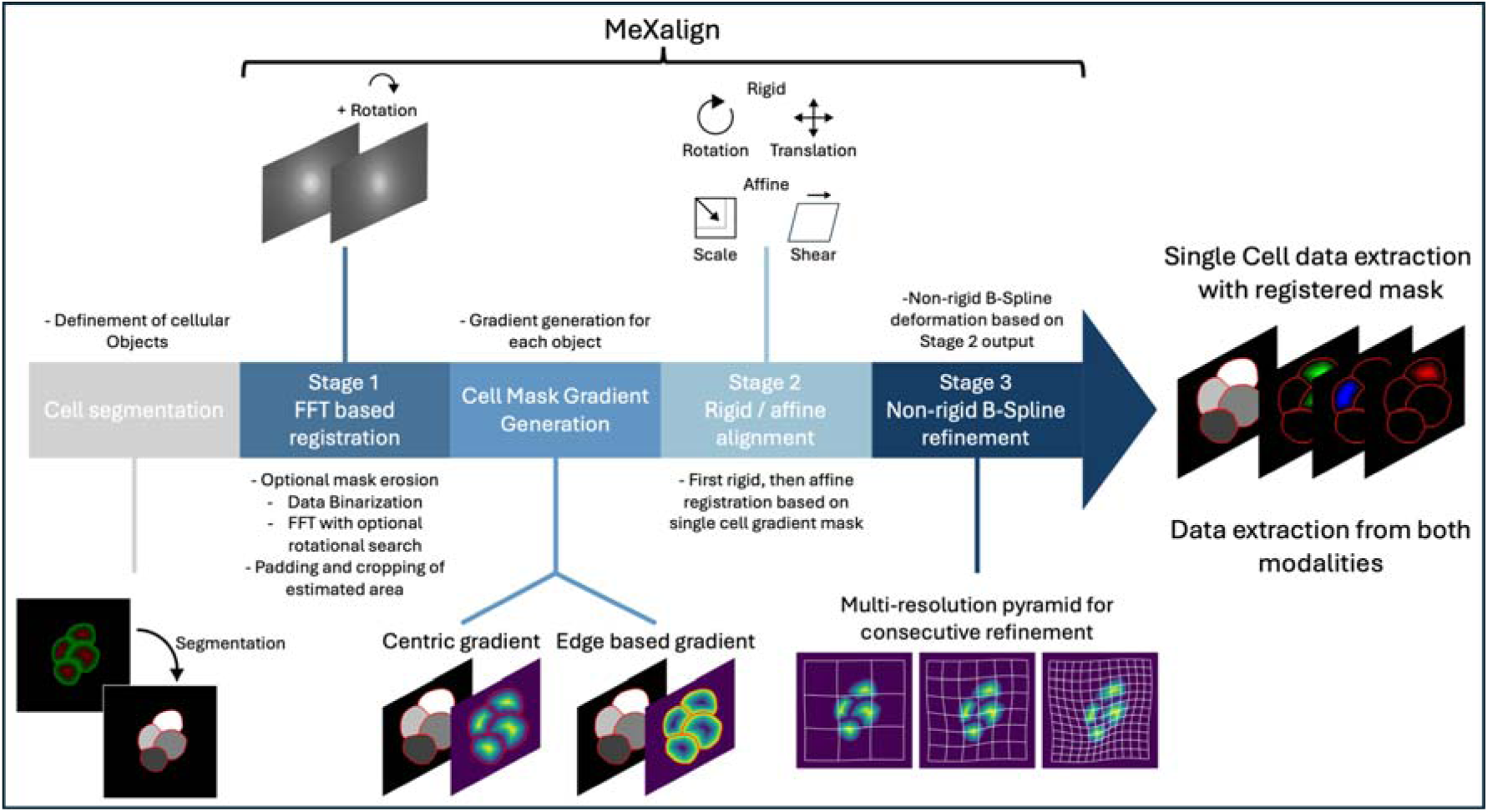
Schematic overview of the MeXalign coregistration pipeline. Following single-cell segmentation, Stage 1 performs FFT-based coarse registration to estimate rotational and translational offsets and identify the corresponding image region. The resulting label masks are converted into cell-based gradient fields, which are subsequently aligned by rigid and affine transformations in Stage 2. In Stage 3, remaining local distortions are corrected by non-rigid B-spline refinement using a multi-resolution approach. The resulting transformations are applied to the selected image channels and label masks, enabling extraction and comparison of registered single-cell data across both imaging modalities.

**Figure 2:**
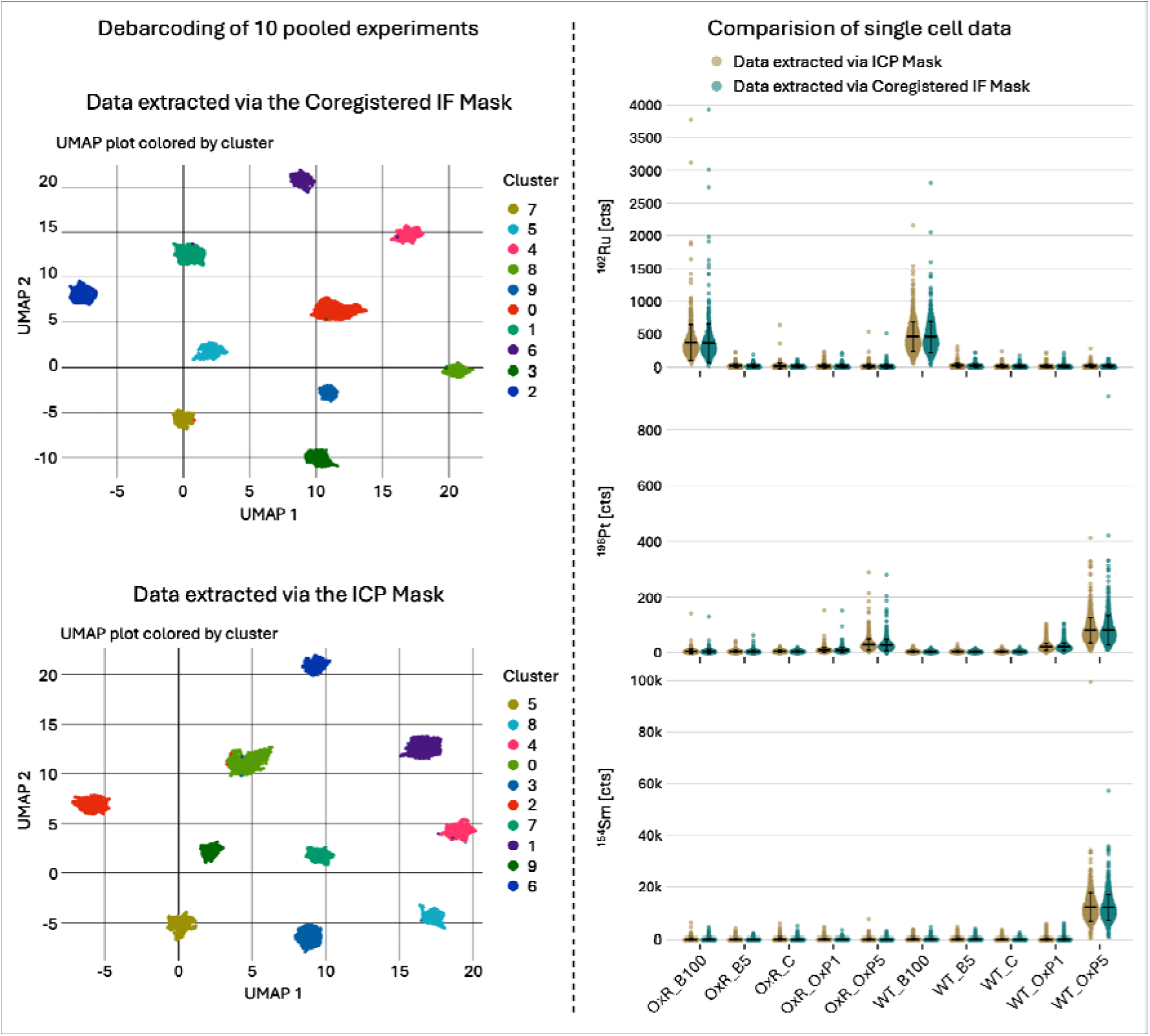
Single cell data comparison between data extracted via an ICP segmentation mask and once via a coregistered IF segmentation mask. **Debarcoding of 10 pooled experiments:** Both UMAPs show ten distinct clusters for ICP and coregistered IF mask derived data. Clustering was performed based on barcode signals using unsupervised Leiden clustering by PhenoGraph. **Comparison of single cell data:** Violin plots of ^102^Ru, ^196^Pt and ^154^Sm (Barcode of WT_OxP5 experiment) across all experiments for ICP derived data (blue) and IF derived data (red) including Mean and standard deviation. Violin plots for the respective experiments show the same distribution and mean. Background signals are equivalent for IF and ICP derived single-cell data.

For WT_B100, single cell data extracted via the ICP mask showed a mean of 454.3 ± 226.4 cts for ^102^Ru and 449.7 ± 239.7 cts for the coregistered IF mask extracted data. For WT_OxP5, single cell data extracted via the ICP mask showed a mean of 81.1 ± 45.4 cts for ^196^Pt and 12145.6 ± 5523.8 cts for ^154^Sm compared to 81.2 ± 53.0 cts for ^196^Pt and 12034.6 ± 4990.6 cts for ^154^Sm for the coregistered IF mask extracted data. All Data show similar results. The close agreement in recorded elemental, barcode and drug signal intensities between data extracted using the native ICP segmentation masks and the coregistered IF-derived masks demonstrates that coregistration preserves single-cell correspondence with sufficient accuracy for quantitative data extraction. This is further supported by the consistent recovery of the ten experimental populations through barcode-based debarcoding and unsupervised clustering. Together, these results show that the calculated transformations accurately transfer the IF-derived cellular masks into the LA-ICP-TOFMS coordinate space without introducing substantial changes to the resulting single-cell measurements. The full list of ^102^Ru, ^196^Pt and barcode signal across all experiments is listed in Table S9.

For the coregistration performance, the herein introduced modified Dice coefficient of 0.885 and IoU of 0.811 show a good overlap of cell masks also after visual verification. The Jacobian determinant remained strictly positive within the cell area of the fixed mask (range 0.825 - 1.197), confirming a locally invertible, fold-free transformation.

#### 3.2.2 Coregistration based on DNA-based segmentation mask and transfer to an alternative IF Membrane segmentation mask

We next tested whether coregistration could be performed using alternative signals for segmentation. Starting from the sample in Figure 2 and Figure S10, we re-segmented it based on the DNA signal for IF (DAPI) and ICP (^193^Ir). This lets us test coregistration on a matched structural signal (DAPI – ^193^Ir, DNA in both modalities).

The coregistration was performed on these DNA derived segmentation masks and the resulting transform was applied to the original WGA-DNA based IF segmentation mask (Input for second optional mask within the GUI). The additional transformed WGA-DNA based IF segmentation mask was then used for data extraction to compare single cell data to the original datasets from the membrane-based segmentations (ICP and IF).

To confirm that coregistration can be performed on DNA and applied to a membrane IF mask, violin plots in Figure S11 compare the three datasets derived from the 10-condition barcoded cytospin sample analyzed above (shown as violin plots of ^102^Ru, ^196^Pt, and the WT_OxP5 barcode signal; means in Table S9). Two serve as references: the original ICP dataset (extracted from the native ICP segmentation) and the standard membrane-based coregistration (membrane-driven, extracted via the IF membrane mask). These are compared against the DNA-based segmentation mask coregistrations. All three are in good agreement, confirming that DNA-driven coregistration recovers the same single-cell data as both the native ICP segmentation and the membrane-based coregistration.

#### 3.2.3 Coregistration based on endogenous elements and transfer to an alternative IF membrane mask

Additionally, a segmentation was performed based on ^23^Na-signal (ICP), a merely similar signal that colocalizes to the DNA. The resulting mask was coregistered towards the DAPI signal (IF) and the resulting transforms applied to the WGA-DNA based IF segmentation mask to extract data and compare single cell data.

A prerequisite for this is that the two signals used for segmentation need to result in single-cell masks and the resulting masks need to roughly co-localize rather than match exactly. The center-based gradients align centers of the masks rather than the exact shape which allows for slightly differing signals to be registerable. If none of these are the case, coregistration is not possible. Best practice is to use dedicated membrane-DNA based segmentation masks, then DNA segmentation mask based coregistration and transfer to a secondary (membrane based) mask and finally on merely similar signal that colocalizes with transfer to a secondary (membrane based) mask.

To confirm that coregistration can be performed on colocalized endogenous elements and applied to a membrane IF mask, violin plots in Figure S11 additionally show the fourth dataset. (shown as violin plots of ^102^Ru, ^196^Pt, and the WT_OxP5 barcode signal; means in Table S9). Data is in good agreement, confirming element-driven coregistration recovers the same single-cell data in this sample compared to native ICP segmentation and the membrane-based coregistration.

#### 3.2.4 Robustness to under-segmentation

The cytospin described in 4.2.1 can be used to validate coregistration performance under progressive under-segmentation. Doublets were generated by merging adjacent cells into single objects. The principle of doublet generation is shown in Figure 3C. This allows defined degrees of under-segmentation to compare coregistration against decreasing segmentation quality. We merged 5, 10, 20, 30, 40 and 50% of cells into doublets. This was achieved by merging adjacent cells to the same label. Coregistration was then performed between the doublet-containing ICP mask and the intact IF mask. To keep the comparison consistent, the registered mask (Figure 3C: Blue outlined mask) was compared against the original ICP mask without doublets (Figure 3C: Red outlined mask). The goal was to test whether the mask registered from a faulty, doublet-containing fixed mask still converges onto the true single-cell layout. If registration degraded with increasing doublets, the shifting doublet boundaries would progressively lower the modified Dice and IoU scores. Instead, both scores remained essentially constant (Figure 3D), confirming that alignment is unaffected by under-segmentation of the fixed mask and overlap of single cells remains steady.

**Figure 3:**
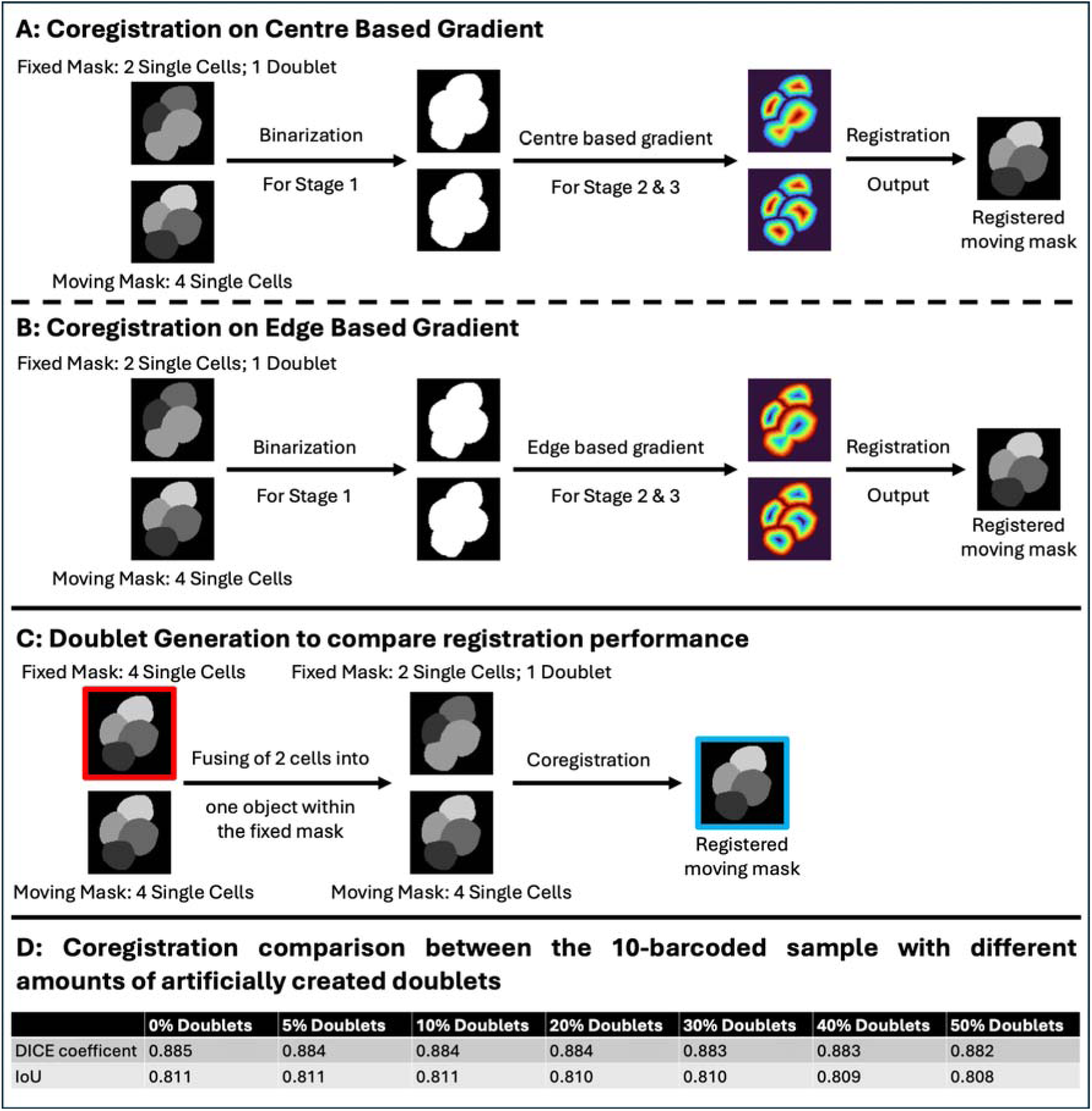
Comparison between 4 cells based on (A) center-based gradient and (B) edge-based gradient with their respective maxima. Within the fixed image, a doublet was created to display the influence on the binarized data (no difference at all) and the gradient fields. This illustrated why convergence on doublets is still accurate. (C) shows the principle of merging two adjacent cells into one doublet (fixed mask) and performing the coregistration. The registered mask (blue) is then compared against the actual fixed mask (red) to calculate the modified Dice coefficient and IoU. (D) shows the modified Dice coefficients and IoU for different number of merged cells (0 % - 50%). Both coefficients are stable if half of cells are merged.

**Figure 4:**
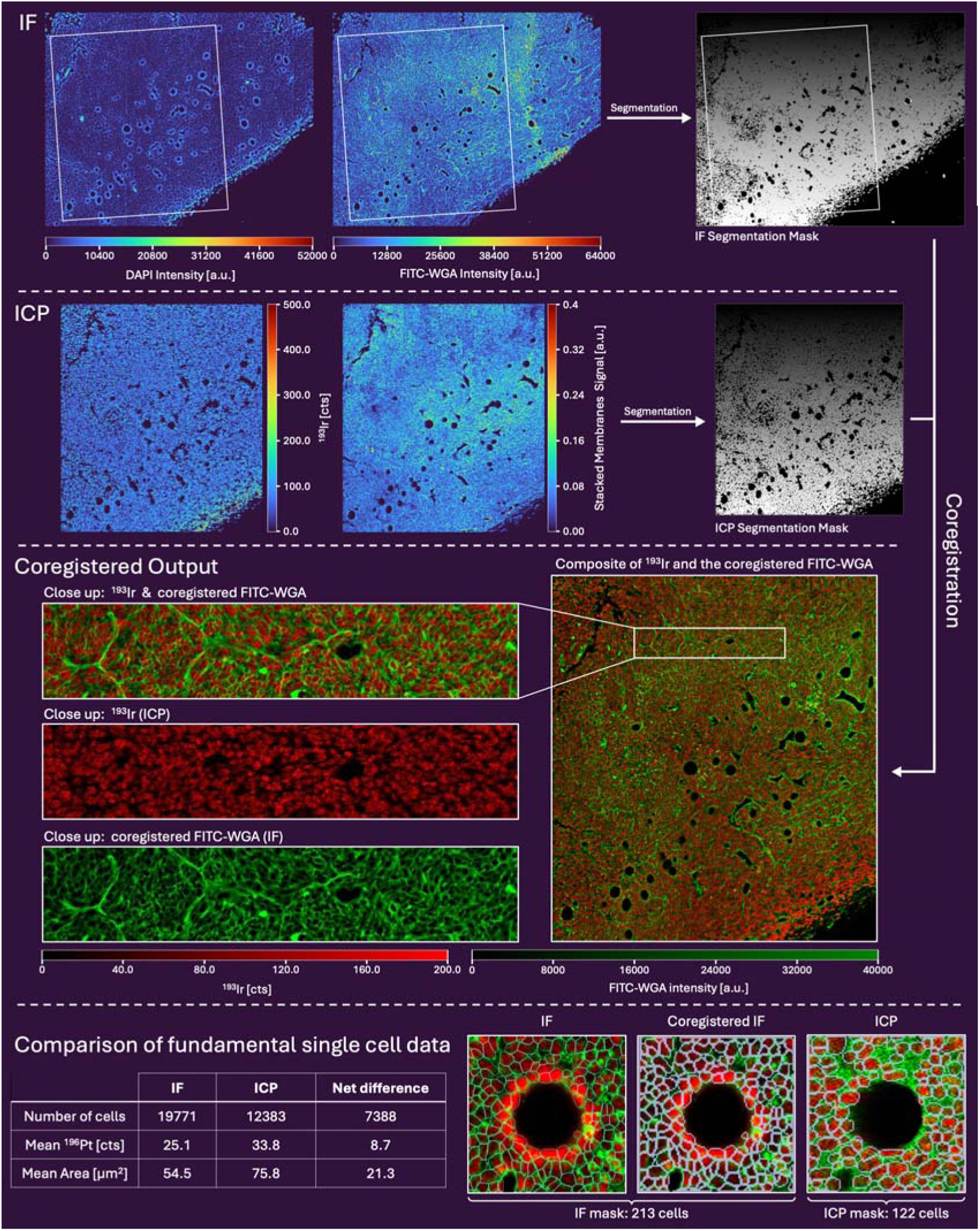
Oxaliplatin treated murine tumor (CT26). Segmentation on both modalities (ICP and IF) was performed using Cellpose-SAM^31^ on the respective DNA and membrane channels with subsequent manual segmentation for remaining unsegmented cells. **IF:** DAPI (DNA) and FITC-WGA (Membrane) was captured via IF. The region corresponds to a larger crop also containing the ablated ICP region (white box). **ICP:** Ablated region of the tumor via LA-ICP-TOFMS. Illustrated are the DNA (^193^Ir-Intercalator) and the membrane channel (Normalized and stacked channels of CD45, beta-Actin, α-SMA, Vimentin, CD44). Coregistration was performed on the respective cell masks. **Coregistered Output:** Illustrated are the composite of the ^193^Ir channel (ICP; red) overlayed with the coregistered FITC-WGA (IF; green) as well as close ups of the composite and the individual signals. **Comparison of fundamental single cell data:** Comparison Between the number of cells mean area of the cells and mean ^196^Pt content per cell. Net difference between ICP and IF is 7388 more cells (+59.7 %), 21.3 µm^2^ less area per cell (−28.1%) and 8.7 less counts of ^196^Pt per cell due to more cells being segmented. On the right-hand side is a close-up view of cells which are undistinguishable within the ICP and the high resolution in IF leads to more cells. IF_Coregistered is the coregistered signal to match the same dimension and coordinates of the ICP image. ICP maps were acquired using 3 µm circular spots with Dosage 3 in X and Y direction resulting in a final pixel size of 1 µm.

The origin of this invariance lies in the per-object gradient representation used for rigid, affine and non-rigid B-Spline stages (Figure 3A & B). Merging two adjacent cells into a single label, whether artificially or through mis-segmentation, does not erase the structure the optimizer relies on, namely the per-object normalized internal distance map. For the center-based gradient, maximum is in the center, minimum at the edges of the object. For a doublet, this produces a bilobed field with two local maxima or an elongated maxima (Figure 3A). Because convergence is intensity driven, and the two or elongated maxima coincide with the two single-cell maxima, the cell centers still align. The major difference is that values between the two cell centers no longer fall to zero as it would happen in two single cells. Moreover, the B-Spline control-point spacing is far coarser than a single cell. The deformation is at each control point driven jointly by many cells in its neighborhood. Hence the two maxima of one doublet are a small contribution among all the correctly one-to-one matched single-cell maxima, further diluting their influence.

The same holds for the edge-based gradient with maxima at the object edges and minimum in the center (Figure 3B). For the doublet, the signal at the internal cell opposition is lost, whereas the outer contours remain the maximum. As the same slide is stained once and then analyzed consecutively, distortions are primarily modality specific distortions and staining difference rather than single cell signals changing. The center-based gradient should nonetheless be more robust when segmentation is imperfect as convergence is towards one maximum in the center, whereas the edge-based gradient presents a closed ring of maxima that can lock onto boundary detail. Additionally, the center-based gradient tolerates modality-specific differences in cell size as the interior maximum stays near the same location even when the border is drawn differently. The edge-based maximum shifts with the contour.

#### 3.2.5 Coregistration based on alternative membrane markers

Within Figure 5, a skin sample was stained using ^153^Eu-labeled Anti-CD44, FITC-WGA and two DNA-intercalators (DAPI and Ir-DNA intercalator) to demonstrate that coregistration can also be performed based on alternative membrane markers signals during segmentation. Segmentation was performed based on the ^153^Eu (Anti-CD44) and ^193^Ir signals for ICP and FITC-WGA and DAPI signals for IF and both masks used for coregistration (Figure S12). LA-ICP-TOFMS segmentation^31^ resulted in 1308 cells whereas for IF in 1405 cells showcasing the improved segmentation within IF To evaluate the coregistration performance, a modified Dice score of 0.874 and an modified IoU of 0.776 were achieved, indicating a robust spatial overlap. The Jacobian determinant remained strictly positive throughout the tissue (range 0.956 - 1.091), confirming a locally invertible, fold free transformation. The primary driver of the structural difference is the marker coverage: CD44 does not stain all cells within skin tissue (limiting the ICP-based segmentation) compared to FITC-WGA (coregistered IF segmentation), which stains the complete tissue extracellular matrix. This comprehensive staining results in larger segmented cell boundaries in areas lacking CD44 expression. Ultimately, this shows that using distinct modality-specific segmentation markers for ICP and IF remains highly compatible and yields highly accurate data. Coregistration parameters are detailed in Table S5.

**Figure 5:**
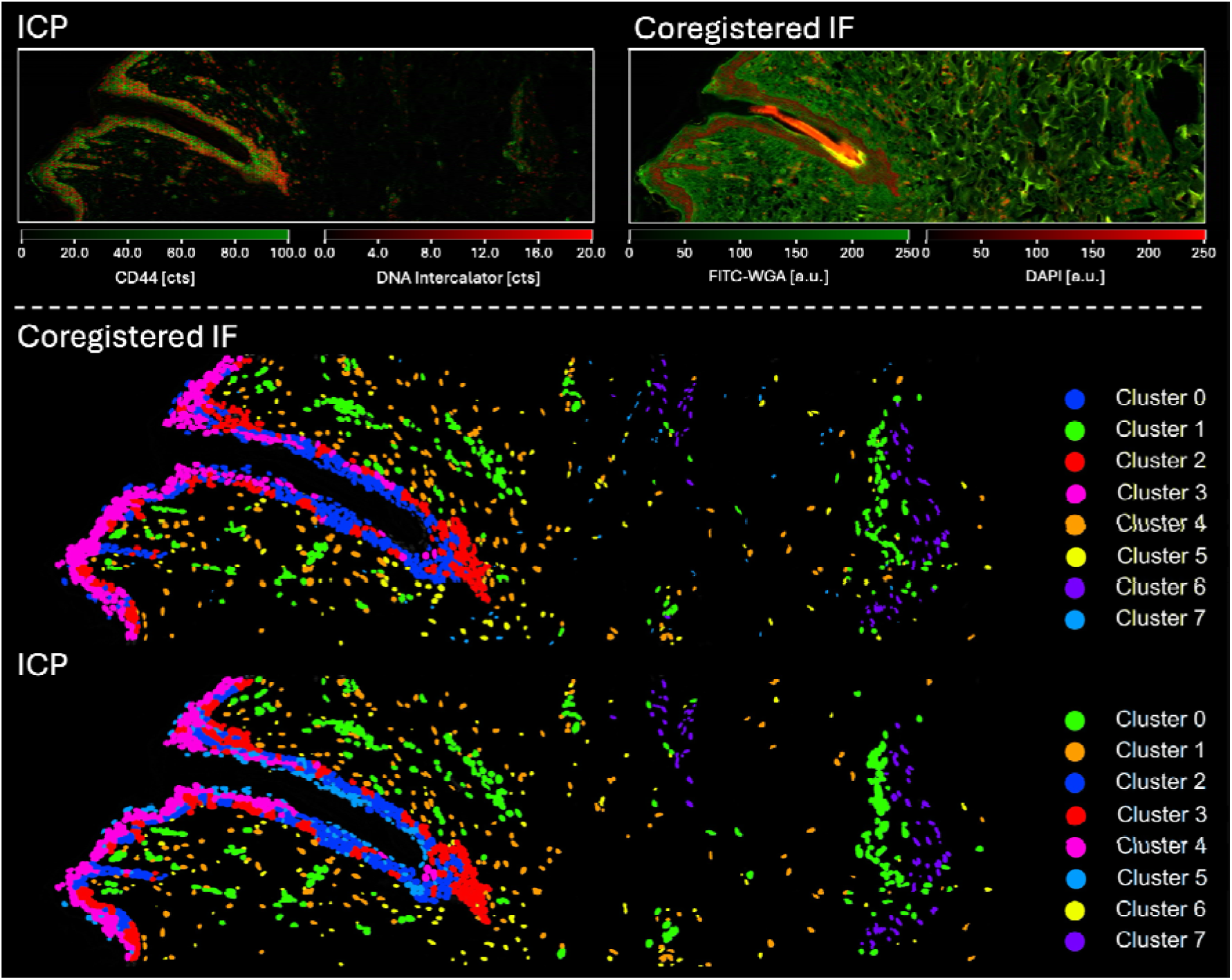
Coregistration between an ex vivo human skin sample analyzed by IF and LA-ICP-TOFMS. Segmentation was performed on the ^193^Ir-DNA-intercalator and CD44 signal for the ICP images and on DAPI and FITC-WGA for IF. After coregistration, clustering was performed based on Vimentin, α-SMA, PanCk, CD34, E-Cad, CD44, Collagen type I and ^56^Fe. For both masks, 8 clusters were obtained. ICP maps were acquired using a 2 µm circular spot with a dosage 4 in the X and Y directions resulting in a final pixel size of 0.5 µm.

#### 3.2.6 Improved segmentation by using higher resolution modalities and achieving quasi subcellular resolution

In IMC, the spatial resolution and sensitivity are inherently coupled. To achieve high-resolution imaging at scales of 250–333 nm^26,27^ results in a quadratic decline in signal intensity per pixel across all channels due to less ablated sample per spot. This is a real limitation, because the principal benefit of high resolution lies precisely in segmentation enabling subcellular and extracellular data extraction.^21,26,27^ Segmentation is further complicated in dense tissues such as tumors, where extreme cellular density and the lack of extracellular matrix hinders the delineation of individual cells.

Coregistration resolves the compromise by decoupling these two requirements. Segmentation accuracy is provided by a high-resolution imaging modality such as IF, while IMC is required at larger spot sizes that retain sufficient signal for quantitative analysis. This way coregistration enables accurate single cell segmentation in IF, transferred to IMC data measured under high sensitivity conditions.

A consequence of insufficient resolution and segmentation is the formation of doublets, the combined assignment of 2 (or more) cells to one segmented object. Doublets bias the statistics as fewer cells are counted resulting in bigger cell diameters as well as increase signals per cell as two or more cells are merged into one cell mask. Here coregistration enables the use of the accurate segmentation mask from the higher resolution modality (IF) to extract IMC data.

Within Figure 4 IF represents the roughly cropped image used for segmentation as well as the IF segmentation mask used for coregistration. The white box corresponds to the via LA-ICP-TOFMS ablated area which is slightly tilted. Segmentation was performed based on the ^193^Ir signal (DNA) and a membrane channel. The latter was generated by individually normalizing and stacking CD45 (^152^Sm), beta-Actin (^154^Sm), α-SMA (^141^Pr), Vimentin (^143^Nd), CD44 (^153^Eu) signals into one layer to enhance cellular and membrane-associated structures. Coregistration parameters are listed in Table S4. Coregistered output represents the coregistered FITC-WGA signal overlayed with the ^193^Ir signal for an intuitive visualization of coregistration performance. Finally single cell data is compared for an Oxaliplatin (9 mg/kg, 24 & 96 h) treated tumor once using the ICP segmentation mask and once the coregistered IF mask for data extraction. The ICP mask shows 12383 single cells compared to the coregistered IF mask with 19771 cells which corresponds to a 59.7 % increase in detected single cells resulting in decrease area per cell by 28.1 % (mean area in ICP: 75.8 µm^2^ vs IF: 54.5 µm^2^). This difference in cell number and mean cell area also affects quantitative single-cell measurements. By resolving previously merged or undetected cells, the measured ICP signal is distributed across a large number of individual cells with smaller areas, which alters the mean signal recorded per cell.

For example, in Figure 4, the mean ^196^Pt signal, representing the oxaliplatin uptake, was 25.7 % lower when extracted using the coregistered IF mask compared to the ICP mask (mean ^196^Pt signal/cell ICP: 33.8 cts vs IF 25.1 cts). The modified Dice coefficient is 0.650 and the modified IoU 0.554. Visual inspection of the overlay confirms good coregistration (Figure 4). The Jacobian determinant remained strictly positive throughout the tissue (range 0.997 - 1.100), confirming a locally invertible, fold free transformation. The slightly reduced Scores compared to the sample within Figure 2 are due to a worse ICP based segmentation as many cells cannot properly be segmented. Hence also the difference of 59.7 % more cells within the coregistered IF mask.

#### 3.2.7 Clustering performance after coregistration

Data extracted from the coregistered IF mask can be used for subsequent single-cell analysis. To demonstrate this application, single cell data was extracted via the ICP mask and registered IF mask (same tissue sections as in Figure S12).

Clustering was performed on single-cell data extracted from both the ICP and coregistered IF masks, using a panel comprising vimentin, α-SMA, pan keratin, CD34, E-cadherin, CD44, collagen type I, and iron (Figure 5). While both modalities revealed a similar distribution of 8 distinct clusters, the IF-derived clusters appeared noticeably more cohesive. The ICP mask tended to over-segment and randomly split the epidermis, whereas the IF mask provided more granular spatial detail. This superior resolution ensured better cell-to-cell separation, which was particularly beneficial for resolving the smaller, densely packed cell populations within the dermis. This illustrates that single cell data retrieved from the coregistered segmentation masks closely align to the actual ICP mask with the benefit of a more accurate segmentation (yielding 7.4% more cells).

#### 3.2.8 Combining LA-ICP-TOFMS and IF data

Beyond overlays and improved segmentation, registration enables signals from both modalities to be combined into a single spatially resolved dataset, thereby increasing readout per sample. In addition to fluorescently labelled antibodies, this provides access to fluorescence-specific readouts that are inaccessible to measurements by LA-ICP-TOFMS including autofluorescence, reporter proteins (e.g. FUCCI cell cycle, senescence, lipid peroxidation, green-fluorescent protein modified cells), subcellular compartments and fluorescent nanoparticles (NPs).

As displayed in Figure 6, fluorescent nanoparticles provide an illustrative example of this complementarity. Since carbon is ubiquitous in biological samples and plastic materials, distinguishing nanoparticle-derived carbon from the endogenous using LA-ICP-MS alone is challenging. The use of fluorescently labelled nanoparticles allows to overcome this limitation. Although LA-ICP-TOFMS enables sensitive elemental detection, its spatial resolution limits the localization of NPs at nanometer scale. Metal doped NPs can facilitate their detection and enables quantification via ICP-TOFMS whereas fluorescence microscopy offers substantially higher resolution for determining their intracellular localization. To demonstrate a potential use case of NP-localization, we want to highlight the spatial accuracy from NP within coregistered images. HCT116 cells were incubated with 490 nm fluorescent polystyrene NPs (Spherotech Sky Blue, 50 ng/mL) and stained using the dual-modality membrane labeling strategy^22,26^ and the corresponding nuclear staining. ICP images were acquired at a resolution of 500 nm (2 µm circular spots with a dosage of 4 in X and Y direction). IF images were acquired at 0.110 µm resolution whereas the IF images within Figure 6 are the registered and transformed images resulting in the 500 nm equivalent. A modified Dice and IoU of 0.887 and 0.823 respectively showed good overlap. No tissue folding was observed (Jacobian determinant 0.969-1.107).

**Figure 6:**
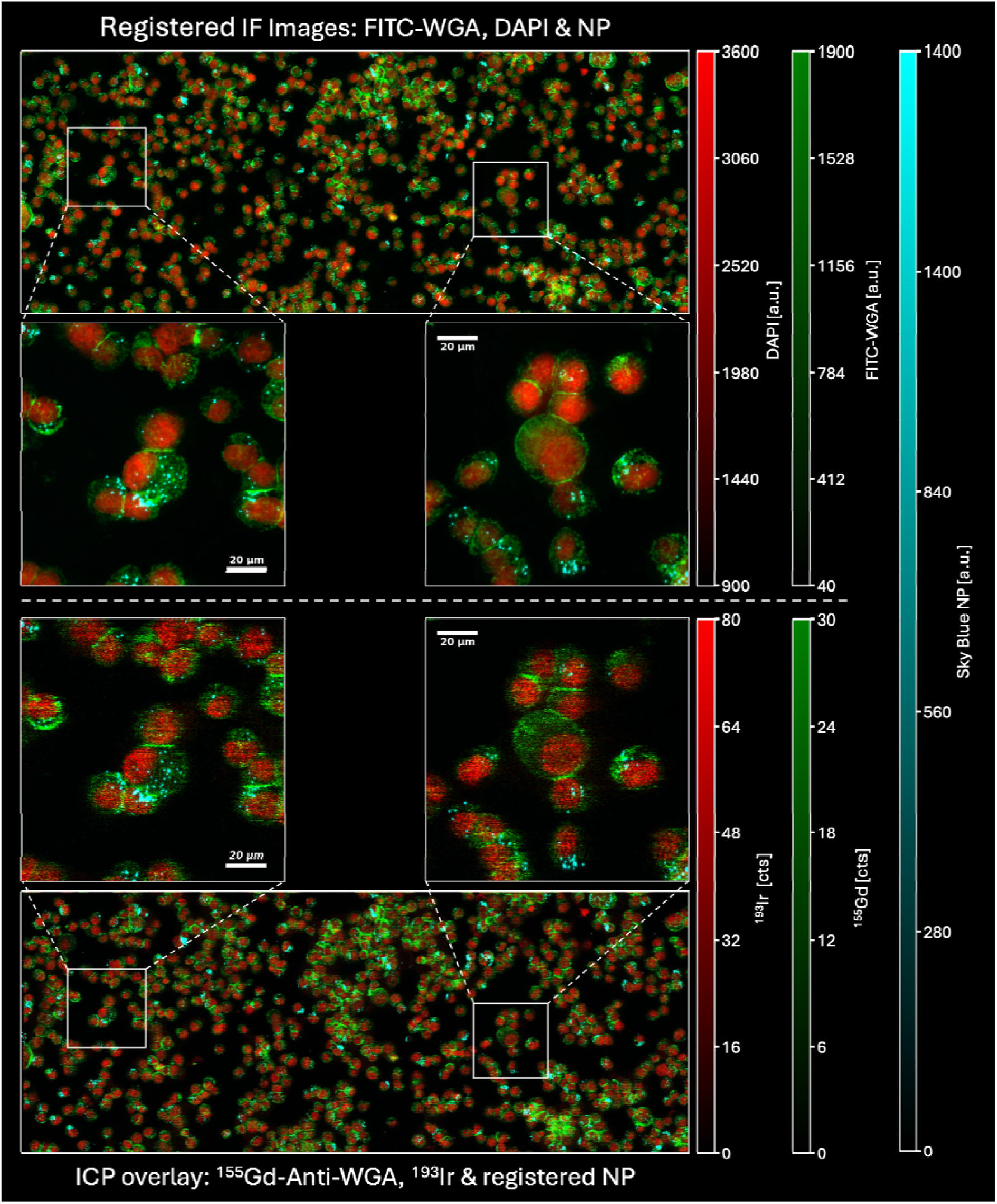
Visualization of coregistration accuracy based on the spatial localization of fluorescent NPs relative to nuclear and membrane staining. HCT116 cells were incubated with 490 nm fluorescent polystyrene NPs (Spherotech Sky Blue, 50 µg/L) for 24h. Images were registered based on DNA-membrane derived segmentation masks. Top: Overlay of the transformed IF images showing the DNA (red), FITC-WGA (green) and SkyBlue PS NPs (cyan). Bottom: corresponding LA-ICP-TOFMS of nuclear signal (red) ^155^Gd-anti-WGA (green) overlaied with the transformed NPs signal (cyan) from the IF dataset. Comparison of the NP signals relative to the nuclear and membrane structure demonstrates preservation of their spatial localization following coregistration and transformation between the two imaging modalities. Magnified regions highlight the same cellular regions of the NP in IF and LA-ICP-TOFMS. ICP maps were acquired using a 2 µm circular spot with a dosage of 4 in the X and Y direction resulting in a final pixel size of 0.5 µm.

Following coregistration (details in Table S7), all three IF channels (DNA, FITC-WGA and NPs) were transformed into the LA-ICP-TOFMS coordinate system (Figure 6). DAPI and the Ir-DNA intercalator and the dual label^22,26^ delineate nuclei and plasma membranes respectively.

When overlaying the transformed NP fluorescence channel with the DNA-intercalator and ^155^Gd-anti-WGA, spatial distribution is preserved from IF to LA-ICP-TOFMS images. This correspondence is particularly evident in the magnified regions, where fluorescent NP features remain localized to the corresponding subcellular regions after registration and transformation into the LA-ICP-TOFMS coordinate system.

#### 3.2.9 Validations of new techniques

Another theoretical use case which was not illustrated within this publication is the use of coregistration as validation of new techniques or combination if data from both modalities. Well-established procedures from one field (e.g. IF) can be fused to a different field (e.g. IMC). Perfect alignment of images is a prerequisite for this.

## 4. Conclusion

MeXalign provides a flexible framework for integrating complementary imaging modalities on the same physical specimen, enabling corresponding cells to be analyzed across modalities while retaining their respective molecular and elemental information. The pipeline was demonstrated in this work using LA-ICP-TOFMS and IF data but is in principle transferable to other imaging modalities that provide sufficient spatial resolution and contrast for cellular segmentation.

Combining FFT-based coarse alignment with rigid/affine and non-rigid B-spline refinement on per-cell gradient fields enabled accurate, fold-free single-cell coregistration, while the quantitative LA-ICP-TOFMS data remained unaltered. Cytospins served as simple model system to first validate the coregistration using the WGA/anti-WGA dual membrane label. This label marks the same structural feature with corresponding signals in both modalities and resulted in modified Dice /IoU scores of 0.885/0.811. The resulting extracted single cell data from the IF derived mask matched the ICP mask. Building on this, coregistration based on alternative signals that colocalize, namely DNA-DNA (^193^Ir/DAPI) or endogenous ^23^Na-DNA (^23^Na/DAPI), with the transform applied to the IF membrane mask, gave equivalent single-cell data. Increasing the complexity from single-cell culture experiments to thin tissue sections, segmentation masks derived from different membrane markers in each modality were also compatible and resulted in a modified Dice/IoU score of 0.874/0.776 for human skin.

The main benefit is that segmentation accuracy and elemental sensitivity are decoupled. In an oxaliplatin-treated murine tumor, IF-derived masks resolved 59.7% more cells than ICP-derived masks (19771 vs 12383), and in skin they gave more cohesive clustering, illustrating improved single-cell detection for downstream analysis. Fluorescent nanoparticles kept their subcellular localization after transfer into the LA-ICP-TOFMS coordinate system, illustrating the readout of complementary information even at subcellular resolution.

Future work should use MeXalign to combine fluorescence-only readouts with quantitative elemental datasets and to validate new techniques against established ones.

## Data Availability

The data supporting the findings of this study are provided in the supplementary material accompanying this article. Upon reasonable request, raw images of the data will be made available. For further inquiries, please contact Gunda Koellensperger at. The NP-dataset to test MeXalign is available at https://doi.org/10.5281/zenodo.24189297.

## Acknowledgements

Open access funding provided by University of Vienna. For open access purposes, the author has applied a CC BY public copyright license to any author accepted manuscript version arising from this submission.

## Supporting Information

Antibody cocktails and LA-ICP-TOFMS parameters, MeXalign GUI parameters and user guide, Coregistration parameters for all samples, Barcoded experiments and mean intensities, Dice and IoU score variants, Visualization of cellular fingerprint by erosion, FFT-based localization of the full cytospin, Barcoded Cytospin Coregistration and single-cell data comparison, Skin coregistration.

