## Supplementary Information for "MeXalign – A Segmentation Mask-Based Registration Pipeline for Multimodal Single-Cell Analysis"

### **Table S1-S****10**

Table S1: Composition of the antibody cocktails for the staining of the tumor and skin sample respectively. All cocktails comprised CD16/32 (diluted 1:100), 5 % BSA in TBS/0.05 % Tween.

| Antibody cocktail for the staining of the tumor | | | |
| --- | --- | --- | --- |
| Antibody target | Clone | Metal Tag | Dilution |
| α-SMA | 1A4 | ^141^Pr | 1:50 |
| Vimentin | D21H3 | ^143^Nd | 1:50 |
| CD45 | D3F8Q | ^152^Sm | 1:50 |
| CD44 | IM7 | ^153^Eu | 1:50 |
| beta-Actin | 2F1-1 | ^154^Sm | 1:50 |
| Anti-WGA |  | ^164^Dy | 1:50 |
| Antibody cocktail for the staining of the skin | | | |
| Antibody target | Clone | Metal Tag | Dilution |
| α -SMA | 1A4 | ^141^Pr | 1:50 |
| Pan-Keratin | C11 | ^148^Nd | 1:50 |
| Vimentin | RV202 | ^149^Sm | 1:50 |
| CD34 | EP373Y | ^151^Eu | 1:50 |
| CD44 | IM7 | ^153^Eu | 1:50 |
| E-Cadherine | 24E10 | ^158^Gd | 1:50 |
| Collagen type I | Polyclonal | ^169^Tm | 1:50 |

Table S2: LA-ICP-TOFMS parameters

| Parameter |  |
| --- | --- |
| ICP-TOFMS |  |
| RF Power [W] | 1440 |
| Sampling depth [mm] | 3.5 |
| Cone materials | Ni |
| Plasma gas flow [L min⁻¹] | 14 |
| Auxiliary gas flow [L min⁻¹] | 0.80 |
| Nebulizer gas flow [L min⁻¹] | 1.06 |
| Measurement mode | Collision cell technology (CCT) |
| CCT gas | 93% He (*ν*/*ν*), 7% H2 (*ν*/*ν*) |
| CCT gas flow [mL min⁻¹] | 4.2 |
| m/z range | 14-256 |
| Laser ablation |  |
| Spot size | 2-3 μm (circular) |
| Interspacing (Y - Overlap) | 0.5 - 1 μm |
| Repetition rate | 300-500 Hz |
| Dosage (X - Overlap) | 2-4 |
| Shot count | 1 |
| Fluence | 0.4 -1.4 J cm⁻² |

Table S3: Tunable parameters within the GUI as well as recommendations and explanation.

| **Parameter (GUI)** | **Applicable within** | **Description and initial** | **Increasing the value typically results in…** | **Decreasing the value typically results in…** |
| --- | --- | --- | --- | --- |
| **Quick settings** | **I**nputs | Quick setting for: MSQ fast: decreased computational workload due to coarser grid.  MSQ high-detail: increased computational workload due to finder grid.  Mattes: Mattes mutual information as metric. |  |  |
| **Mask of Fixed Image (Reference)** | Inputs | Selects the fixed reference mask defining the target coordinate system. | / | / |
| **Mask of Moving Image** | Inputs | Selects the moving mask that is transformed to match the fixed mask. | / | / |
| **Fixed Image Pixel size (µm/pixel)** | Inputs | Insert the pixel size of the fixed image. | / | / |
| **Moving pixel size (µm/px)** | Inputs | Insert the pixel size of the moving image. | / | / |
| **Gradient generation** | Inputs | Center based: Cell cores are maximum, cell edges minimum  Edge based: Edges are maximum, cell cores minimum  Applies to Rigid/Affine & B-Spline | Center-based is more stable especially when Segmentation differs. Convergence is based on a single point (Center) per cell.  -  The edge-based can be used when the segmentation is very similar. Convergence is based on a ring of maxima at the edges of each object. | When segmentation differs too much or there are too many missing cells, Edge based gradient can create artefacts as convergence based on adjacent missing cells will be overoptimized. |
| **Second mask (Optional)** | Inputs | A second label mask warped with nearest neighbour (preserves cell IDs) | / | / |
| **Erosion Radius (px)** | FFT | Tick box to enable erosion.  Pixels removed from each object boundary before label mask binarization. Used to create inter-object gaps (“cellular fingerprint”). Erosion is internally scaled between modalities based on pixel size.  Recommendation: 1-2 pixels for dense samples. | More separation between adjacent objects and including more background. This improves distinctness in dense samples. Excessive erosion can remove informative structure and weaken matching. | Preserves object geometry. |
| **Crop Padding (px)** | FFT | Padding around the auto-cropped ROI. (to reduce data during Rigid/Affine and B-Spline) | Increase if image is too much rotated. Better would be to manually rotate the IF images before registration. | / |
| **Enable rotation search + Rotation range (±°) + step size (°)** | **FFT** | **The rotation pre-alignment sweep (Default: ±3° range with 0,25° steps)** | **Increase if rotation between images is bigger. For bigger differences, manually rotate images before registration. Major rotations can lead to cut iff part of the matching region. In this case, padding needs to be increased.** | **/** |
| **Number of iterations** | Rigid/Affine  B-Spline | Maximum number of optimizer update steps per stage.  Automatically stops iteration when reaching a plateau. | Greater chance of convergence. Can improve alignment but increases runtime. | Faster runtime however at risk of premature stopping (underfitting), leaving residual misalignment for later stages. |
| **Learning rate** | Rigid/Affine | Optimizer step size controlling how aggressively parameters are updated.  Recommendation: 1 for small /medium images and 0.5 for bigger images | Faster convergence and larger updates. This can correct larger mismatches but increases the risk of overshooting. | More stable fine refinement however slower convergence. |
| **Metric sampling rate** | Rigid/Affine | Fraction of pixels sampled to compute the similarity metric each iteration. | More stable/reproducible metric; often improves fine-stage stability; slower computation per iteration. | Faster computation but potential worsens convergence. |
| **Shrink factors (comma-separated list)** | Rigid/Affine | Multi-resolution downsampling schedule from coarse to fine (e.g., 4,2,1). Controls capture range and stability of optimization. Always from high to low. Recommendation: 4,2,1  For Rigid it is hardcoded as 8,4,2 | More aggressive early coarsening (larger initial factors): greater capture range and stability for large deformations; delays fine-detail matching until later.  The larger the images get, the more distortion can appear. For extremely large images, we recommend increasing the amount of steps to e.g.,[16,8,4,2,1] or even [64,32,16,8,4,2,1] | Less coarsening: earlier fine-detail influence; reduced capture range and potentially less stable convergence if residual misalignment is large. |
| **Metric** | B-Spline | MSQ (mean squares): The mean of the squared pixel-intensity differences between the fixed and moving gradient images, evaluated within the fixed-cell mask; minimized during optimization.  Mattes: Mattes Mutual Information. A Parzen-window (histogram) estimate of the mutual information between the two intensity distributions, maximized during optimization. | / | / |
| **Control Point Spacing (µm)** | B-Spline | Defines the number of B-spline control-point intervals. Image Dimension (in px) are multiplied by pixel size (in µm/px) to get physical dimensions. Control points are set every 150 µm.  This will form the grid. | Increasing control point spacing will lead to a coarser mesh.  Coarser mesh: smoother and more regular deformation. Faster runtime and less overfitting but at cost of less local distortions. | Decreasing control point spacing will lead to a finer mesh.  Finer mesh (more control points): more local flexibility and better correction of local distortions. However slower and increased risk of overfitting to mask defects. |
| **Mesh scale factor**^36,46,47^ **(comma-separated list)** | B-Spline | B-Spline undergoes several run with finer grids resulting in time saving and better overlap. This needs to be in accordance with the with **Shrink factors.**  e.g., Mesh scale factor 1,2,4 => Shrink factor 4,2,1  or Mesh scale factor 1,3,6,9 => Shrink factor 9,6,3,1 | More B-Spline runs lead to increase running time but potentially better convergence  For large images, distortions get proportionally bigger. Here adding B-Spline runs, e.g., Mesh scale factors 1,2,4,8 and Shrink factors 8,4,2,1 | Less B-Spline runs lead to faster running time but potentially worse convergence. Especially for major distortions.  Coarse grids may lead to worsen convergence. |
| **Shrink factors (comma-separated list)** | B-Spline | Multi-resolution downsampling schedule from coarse to fine (e.g., 4,2,1). Controls capture range and stability of optimization. Always from high to low.  Recommendation: 4,2,1 | More aggressive early coarsening (larger initial factors): greater capture range and stability for large deformations; delays fine-detail matching until later. | Less coarsening: earlier fine-detail influence; reduced capture range and potentially less stable convergence if residual misalignment is large. |
| **Early-stop window** | B-Spline | Number of iterations back the metric is compared; 0 disables. Default 3. | / | / |
| **Early-stop tolerance** | B-Spline | Min relative metric improvement over that window, below it will stop further iterations | / | / |

Table S4: Coregistration parameters used for the Tumor sample to compare improved segmentation.

| **Parameter** | **FFT** | **Rigid/Affine** | **B-Spline** |
| --- | --- | --- | --- |
| **R**otation Range | 3.00° | **/** | **/** |
| **Rotation Step Size** | 0.25° | **/** | **/** |
| **Erosion** | Yes | **/** | / |
| **Number of Iterations** | / | Coarse 100 / Affine 200 | 200 |
| **Learning Rate** | / | Coarse 1.0 / Affine 0.5 | / |
| **Metric Sampling Rate** | / | Coarse 1.0 / Affine 1.0 | / |
| **Shrink factors** | / | Affine 4,2,1 | 4,2,1 |
| **Mesh Scale Factor** | / | / | 1,2,4 |
| **Initial Grid** | / | / | 6,6 |
| **Metric** | / | / | mattes |

Table S5: Coregistration parameters used for the Skin sample to illustrate comparable clustering.

| **Parameter** | **FFT** | **Rigid/Affine** | **B-Spline** |
| --- | --- | --- | --- |
| **R**otation Range | 3.00° | **/** | **/** |
| **Rotation Step Size** | 0.25° | **/** | **/** |
| **Erosion** | No | **/** | / |
| **Number of Iterations** | / | Coarse 100 / Affine 200 | 40 |
| **Learning Rate** | / | Coarse 1.0 / Affine 0.5 | / |
| **Metric Sampling Rate** | / | Coarse 1.0 / Affine 1.0 | / |
| **Shrink factors** | / | Affine 4,2,1 | 4,2 |
| **Mesh Scale Factor** | / | / | 1,2 |
| **Initial Grid** | / | / | 15,5 |
| **Metric** | / | / | msq |

Table S6: Coregistration parameters used for the 10 experiments pooled cytospin sample to compare debarcoding and single cell data extracted from both masks.

| **Parameter** | **FFT** | **Rigid/Affine** | **B-Spline** |
| --- | --- | --- | --- |
| **R**otation Range | 3.00° | **/** | **/** |
| **Rotation Step Size** | 0.25° | **/** | **/** |
| **Erosion applied** | No | **/** | / |
| **Number of Iterations** | / | Coarse 100 / Affine 200 | 20 |
| **Learning Rate** | / | Coarse 1.0 / Affine 0.5 | / |
| **Metric Sampling Rate** | / | Coarse 1.0 / Affine 1.0 | / |
| **Shrink factors** | / | Coarse 8,4,2 / Affine 4,2,1 | 4,2 |
| **Mesh Scale Factor** | / | / | 1,2 |
| **Initial Grid** | / | / | 24,8 |
| **Metric** | / | / | msq |

Table S7: Coregistration parameters used for the HCT116 cells treated with 490 nm polystyrene NPs (Spherotech sky Blue, 50 µM, 24h) cytospin sample to illustrate registration accuracy.

| **Parameter** | **FFT** | **Rigid/Affine** | **B-Spline** |
| --- | --- | --- | --- |
| **R**otation Range | 3.00° | **/** | **/** |
| **Rotation Step Size** | 0.25° | **/** | **/** |
| **Erosion applied** | No | **/** | / |
| **Number of Iterations** | / | Coarse 100 / Affine 200 | 40 |
| **Learning Rate** | / | Coarse 1.0 / Affine 0.5 | / |
| **Metric Sampling Rate** | / | Coarse 1.0 / Affine 1.0 | / |
| **Shrink factors** | / | Coarse 8,4,2 / Affine 4,2,1 | 4,2,1 |
| **Mesh Scale Factor** | / | / | 1,2,4 |
| **Initial Grid** | / | / | 12,5 |
| **Metric** | / | / | MMI |

Table S8: Abbreviations for the ten barcoded and pooled experiments

|  | Treatments | | | | |
| --- | --- | --- | --- | --- | --- |
|  | Control | OxP 1µM | OxP 5µM | BOLD-100 5µM | BOLD-100 100µM |
| HCT116 WT  Barcode: | WT_C  ^143^Nd | WT_OxP1  ^144^Nd | WT_OxP5  ^154^Sm | WT_B5  ^147^Sm | WT_B100  ^149^Sm |
| HCT116 OxR  Barcode | OxR_C  ^151^Eu | OxR _OxP1  ^152^Sm | OxR _OxP5  ^145^Nd | OxR _B5  ^155^Gd | OxR _B100  ^165^Ho |

Table S9: Mean intensities across ten pooled and debarcoded experiments.

|  | **Intensities of signals extracted via the ICP mask** | | | **Coregistration via Membrane based segmentation masks.**  **Intensities of signals extracted via the coregistered IF mask** | | |
| --- | --- | --- | --- | --- | --- | --- |
|  | **Mean ^102^Ru/cell [cts]** | **Mean ^196^Pt/cell [cts]** | **Mean WT_OxP5 barcode signal/cell [cts]** | **Mean ^102^Ru/cell [cts]** | **Mean ^196^Pt/cell [cts]** | **Mean WT_OxP5 barcode signal/cell [cts]** |
| OxR_B100 | 363.8 ± 276.8 | 5.7 ± 6.9 | 204.3 ± 441.2 | 354.8 ± 295.5 | 5.4 ± 6.3 | 185.5 ± 389.9 |
| OxR_B5 | 19.2 ± 20.8 | 5.3 ± 3.8 | 188.0 ± 363.2 | 17.0 ± 16.6 | 4.9 ± 4.1 | 149.6 ± 249.9 |
| OxR_Ctr | 17.1 ± 35.3 | 5.9 ± 3.6 | 227.1 ± 495.7 | 14.0 ± 15.8 | 5.3 ± 3.1 | 178.2 ± 378.8 |
| OxR_OxP1 | 15.4 ± 24.4 | 10.4 ± 8.5 | 219.4 ± 367.0 | 13.3 ± 19.3 | 10.0 ± 8.8 | 189.2 ± 226.9 |
| OxR_OxP5 | 13.8 ± 24.7 | 29.8 ± 20.0 | 205.6 ± 425.0 | 12.3 ± 22.1 | 28.6 ± 19.9 | 159.1 ± 269.3 |
| WT_B100 | 454.3 ± 226.4 | 5.0 ± 3.3 | 249.6 ± 431.5 | 449.7 ± 239.7 | 4.7 ± 3.0 | 218.5 ± 370.7 |
| WT_B5 | 20.6 ± 25.5 | 5.0 ± 3.3 | 247.1 ± 453.1 | 18.8 ± 19.8 | 4.6 ± 2.8 | 218.4 ± 362.0 |
| WT_Ctr | 13.4 ± 19.0 | 5.0 ± 3.1 | 198.3 ± 345.6 | 12.1 ± 16.1 | 4.8 ± 2.9 | 170.5 ± 258.0 |
| WT_OxP1 | 11.9 ± 16.6 | 21.4 ± 12.1 | 195.1 ± 452.3 | 11.5 ± 15.4 | 21.8 ± 12.9 | 191.3 ± 453.7 |
| WT_OxP5 | 16.1 ± 21.2 | 81.1 ± 45.4 | 12145.6 ± 5523.8 | 15.2 ± 17.8 | 81.2 ± 53.0 | 12034.6 ± 4990.6 |
|  | **Coregistration via DNA based segmentation masks (ICP and IF).**  **Extraction via transformed IF membrane segmentation mask** | | | **Coregistration via Na (ICP) and DNA (IF) based segmentation masks.**  **Extraction via transformed IF membrane segmentation mask** | | |
|  | **Mean ^102^Ru/cell [cts]** | **Mean ^196^Pt/cell [cts]** | **Mean WT_OxP5 barcode signal/cell [cts]** | **Mean ^102^Ru/cell [cts]** | **Mean ^196^Pt/cell [cts]** | **Mean WT_OxP5 barcode signal/cell [cts]** |
| OxR_B100 | 354.6 ± 294.5 | 5.4 ± 6.7 | 174.7 ± 315.7 | 353.0 ± 293.5 | 5.4 ± 6.9 | 172.6 ± 322.1 |
| OxR_B5 | 17.1 ± 16.9 | 4.9 ± 4.1 | 148.7 ± 252.4 | 17.1 ± 16.8 | 4.9 ± 3.6 | 153.8 ± 267.2 |
| OxR_Ctr | 14.1 ± 15.8 | 5.3 ± 3.1 | 179.2 ± 370.8 | 13.9 ± 15.5 | 5.2 ± 3.1 | 173.8 ± 339.7 |
| OxR_OxP1 | 13.3 ± 19.2 | 10.0 ± 8.8 | 195.6 ± 253.7 | 13.3 ± 19.0 | 10.0 ± 8.8 | 193.5 ± 243.7 |
| OxR_OxP5 | 11.8 ± 15.7 | 28.8 ± 19.9 | 170.4 ± 337.1 | 11.8 ± 16.0 | 28.7 ± 19.8 | 175.8 ± 391.0 |
| WT_B100 | 449.3 ± 239.4 | 4.6 ± 2.9 | 214.9 ± 369.0 | 447.8 ± 240.0 | 4.7 ± 3.0 | 203.8 ± 312.4 |
| WT_B5 | 18.4 ± 18.1 | 4.6 ± 2.8 | 214.7 ± 357.5 | 19.0 ± 20.9 | 4.7 ± 3.0 | 217.8 ± 365.2 |
| WT_Ctr | 11.8 ± 14.7 | 4.8 ± 2.9 | 171.9 ± 269.1 | 11.9 ± 14.9 | 4.8 ± 2.9 | 178.2 ± 296.8 |
| WT_OxP1 | 11.5 ± 15.3 | 21.8 ± 12.8 | 185.2 ± 431.7 | 11.4 ± 15.0 | 21.7 ± 12.7 | 186.5 ± 443.9 |
| WT_OxP5 | 15.2 ± 17.7 | 81.2 ± 53.2 | 12000.6 ± 5010.2 | 15.6 ± 19.7 | 80.8 ± 53.3 | 11935.5 ± 5030.5 |

Table S10: Coregistration parameters used for the complete Cytospin sample to illustrate the FFT finding the correct spot on the sample.

| **Parameter** | **FFT** | **Rigid/Affine** | **B-Spline** |
| --- | --- | --- | --- |
| **R**otation Range | 3.00° | **/** | **/** |
| **Rotation Step Size** | 0.25° | **/** | **/** |
| **Erosion** | No | **/** | / |
| **Number of Iterations** | / | Coarse 100 / Affine 200 | 200 |
| **Learning Rate** | / | Coarse 1.0 / Affine 0.5 | / |
| **Metric Sampling Rate** | / | Coarse 1.0 / Affine 1.0 | / |
| **Shrink factors** | / | Affine 4,2,1 | 4,2,1 |
| **Mesh Scale Factor** | / | / | 1,2,4 |
| **Initial Grid** | / | / | 6,6 |
| **Metric** | / | / | mattes |

### **Figures S1 – S12**

**The derivation of the modified Dice coefficient and IoU to compare segmentation/coregistration performance**

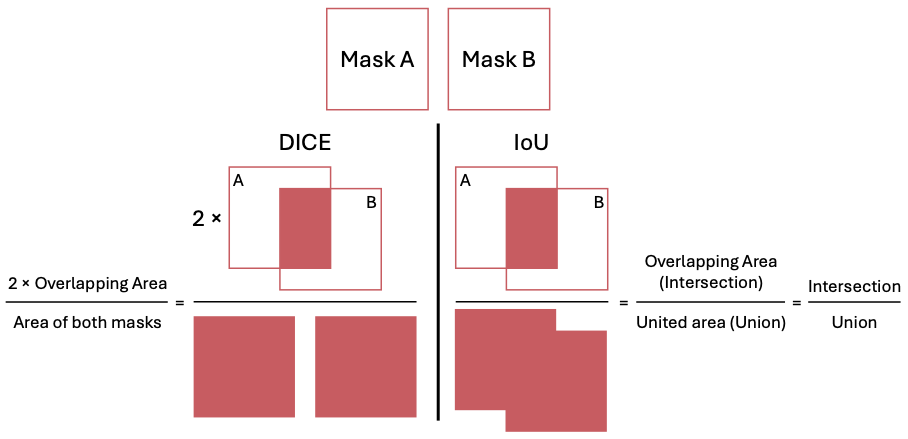

Figure S1: Visual explanation of the Dice and IoU score. The Dice score consists of twice the overlapping area of both masks divided by the sum of both masks. The Intersection of Union (IoU) consists of the overlapping area of both masks (intersection) divided by the united overlapping area (Union).

Two common validation metrics used for comparing mask, tissue or segmentation masks are the Dice coefficient and Intersection over Union (IoU, also named Jaccard Index). The Dice coefficient is calculated as twice the overlapping area of two masks divided by the sum of their individual areas. The IoU is calculated as the overlapping area of two masks (intersection) divided by the combined area (Union) (see Figure S1). Both range from 0 (no overlap) to 1 (identical masks).

However, these metrics come with caveats because one of the major purposes of a coregistration here is the improved segmentation.^6,7^ Improved segmentation means being able to segment cells which previously could not be segmented or the reduction of multiplets, so two or more cells which could not be differentiated and were segmented as one cell even though they consisted of multiple cells. For this reason, we present five complementary Dice and IoU scores.

Mask A contains 3 objects: 2 single cells (A1 and A2) and 1 doublet (A3), so two cells being segmented as one. This represents the low-resolution image.

Mask B Contains 5 objects: 5 single cells (B1 – B5) with slightly different segmentation outlines compared to mask A. In this mask, the doublet within Mask A was successfully segmented as two individual cells (B1 and B4). One additional cell was segmented (B3). This represents the high-resolution image with improved segmentation.

The different scores possible:

1. Silhouette based score: Dice coefficent and IoU are calculated based on binarized masks (any cell = foreground = 1; background = 0). A silhouette-based score evaluates global tissue alignment. All objects contribute but cell identity is lost. So, no statement about cell segmentation quality can be made. A coregistration can achieve a high silhouette score while misalignments at cellular level are not considered. For this, the following object-level scores should be considered.

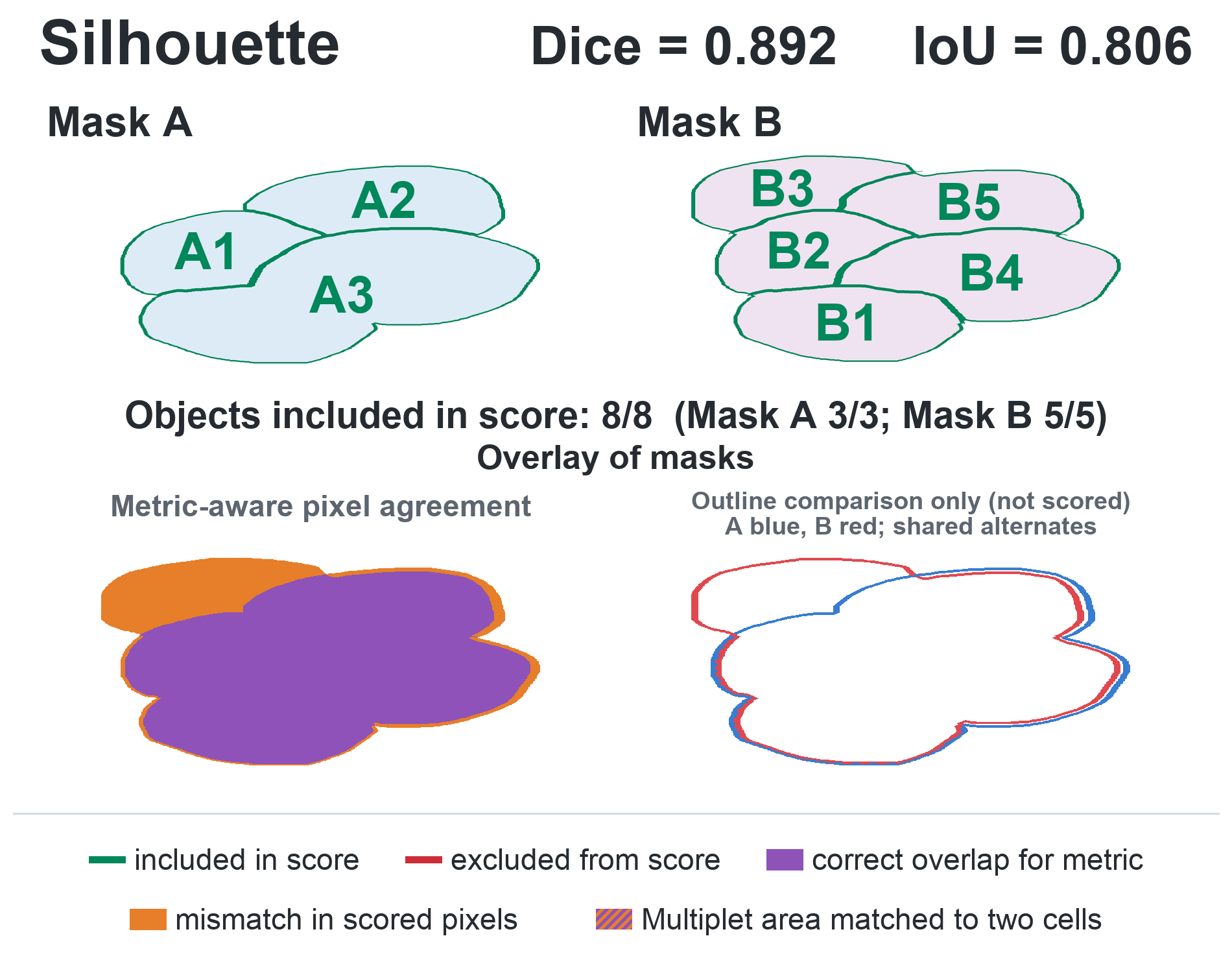

Figure S2: Silhouette based scores.

Scores 2-5 are evaluated per object/cell rather than on the binarized silhouette. This is more informative about cell-to-cell overlap and let us assess whether individual cells actually overlap, rather than just the global structure. They use a fixed correspondence criterion. **Two cells are considered as match when their shared (overlapping) area covers at least 50% of the area of at least one of the two cells.** The Dice coefficient and IoU reported are the mean of the individual object-level scores.

2. All-cell score: Every cell in both masks is independently compared to a respective counterpart. If that overlap exceeds 50 of at least one of the two cells, it is counted as a match. If a cell has no matching counterpart, it is still included with a score of 0 in the mean. Because the comparison runs in both directions, this score penalizes multiplets. When two cells are both matched to one doublet (B1 => A3 and B4 => A3), each comparison is scored separately. Hence the doublet area is effectively counted more than once. This artificially increases the area and lowers the score as the doublet area is incorporated twice for A3. Once for B1 => A3 and once for B4 => A3.

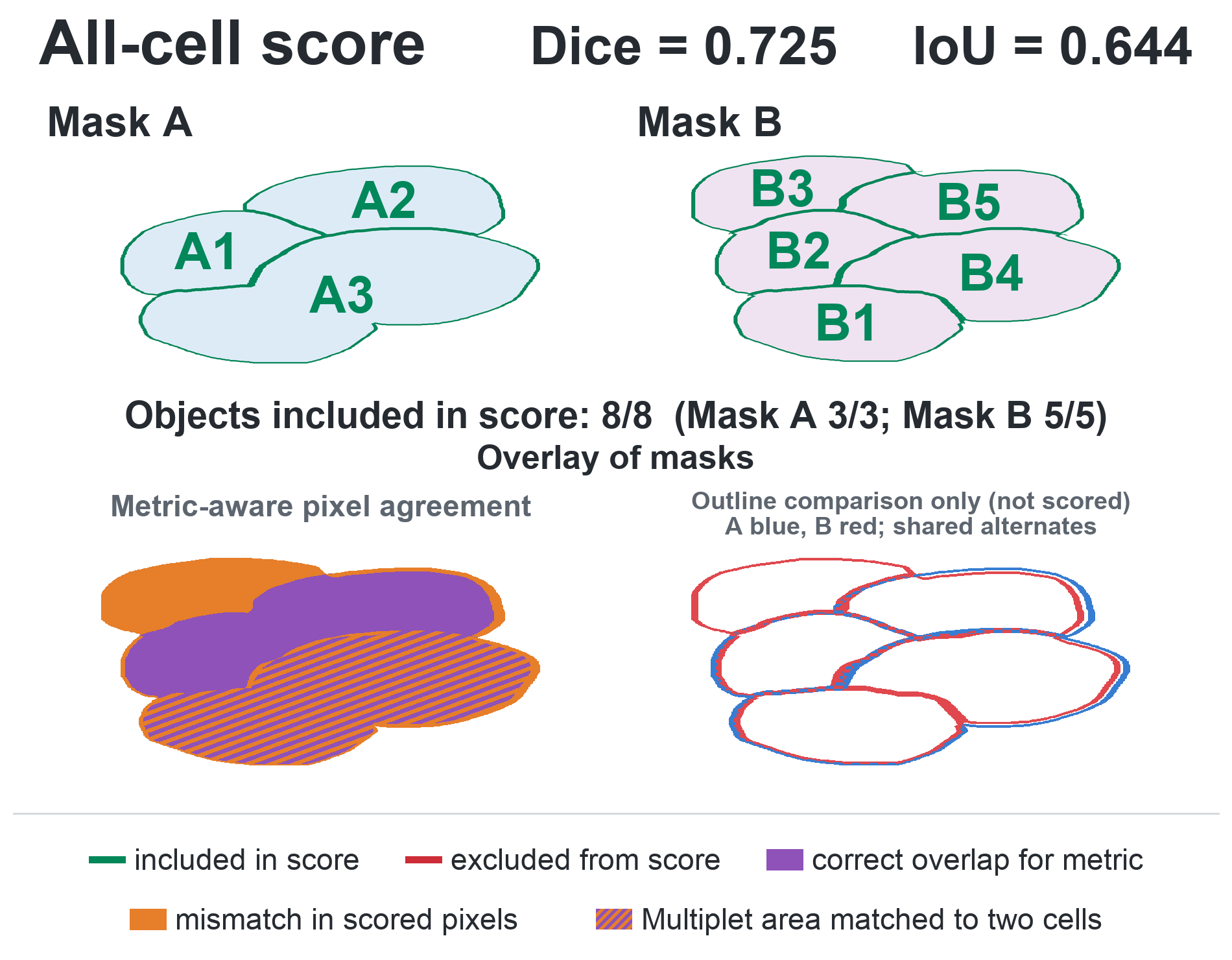

Figure S3: All-cell based Score.

3. Only 1:1 Matches: Only cells which have exactly one-to-one correspondence (one A cell to one B cell) contribute to the final score. Unmatched cells (e.g. B3) or doubly matched cells (A3 and with this the corresponding B1 and B4). This serves as alignment quality of true matched cells. This score is overoptimistic as mismatches are excluded and serves more as comparison for cell matching. This reflects alignment quality of unambiguously matched cells. It is the most optimistic of the object-level-scores because the cells where segmentation differs are excluded. With this score, the number of unmatched cells and doublets not considered should be stated.

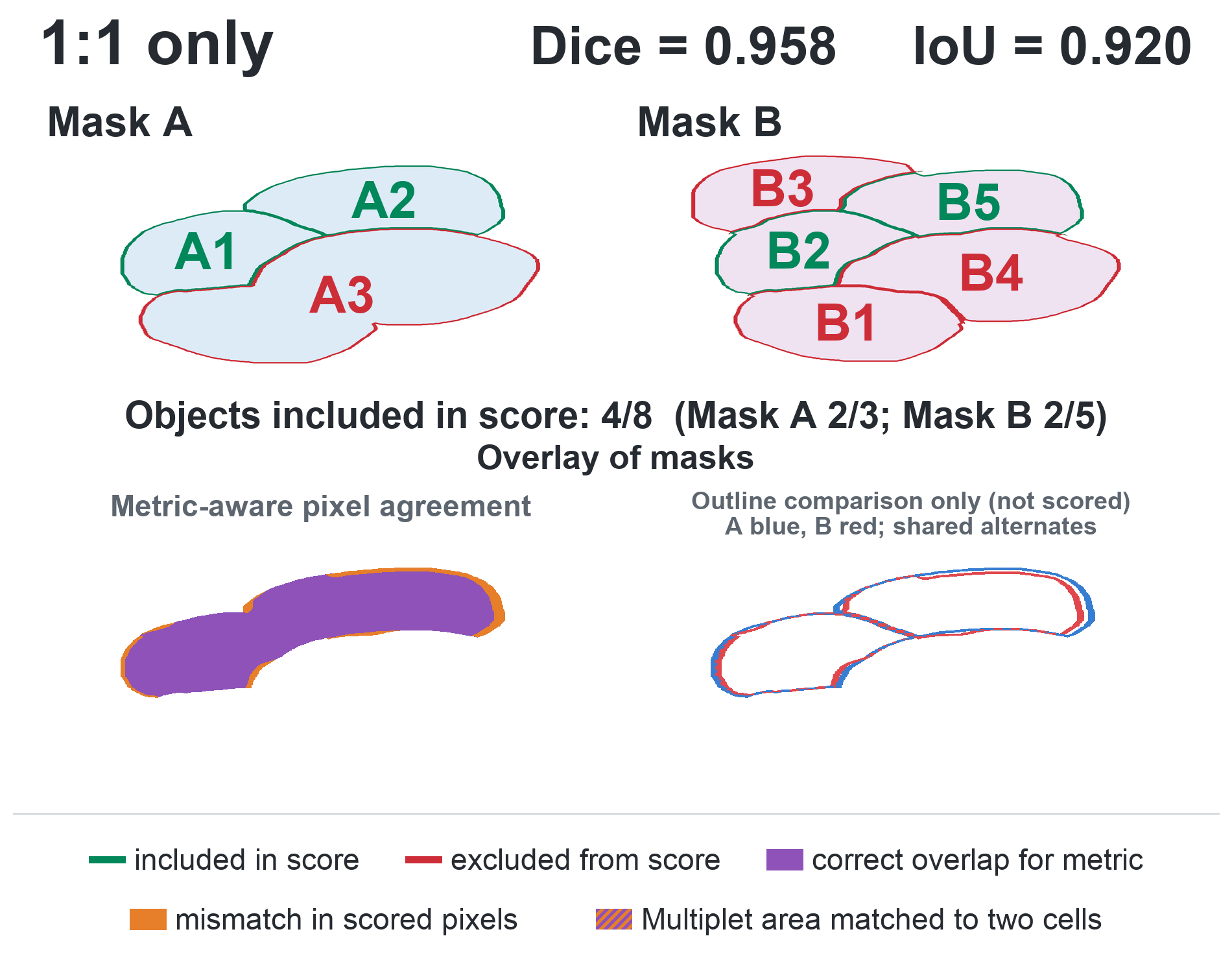

Figure S4: One-to-one only match based scores.

4. 1:1 and fused cell groups: This score includes the same matches as the previous score (Cell A1 => B1 and A2 => B5) but also implement multiplets. The two single cells (B1 & B4) which are matched to the doublet (A3) are fused into one mask and compared to A3 as a single unit. This removes the double counting present in score 2, however slightly overestimates coregistration performance, because the fused B1+B4 area is compared against the doublet A3 as one combined region. Segmentation accuracy of the border between B1 and B4 is not assessed, since the doublet A3 is a single object and provides no internal border to compare against.

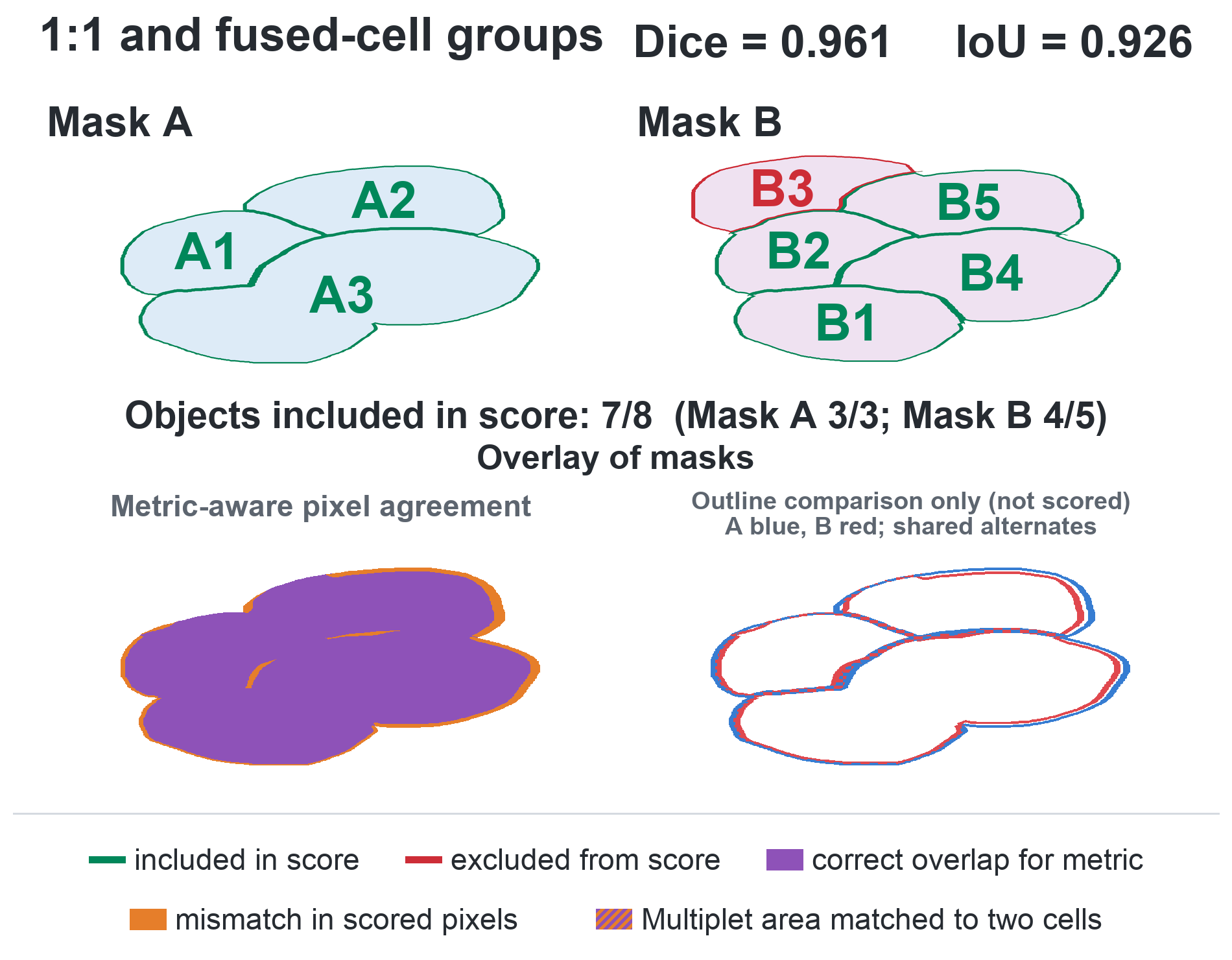

Figure S5: One-to-one only match + fused-cell groups based scores.

5. Complete / Modified score.

The modified score combined the other scores. It uses the 1:1 matched cells (A1 => B1 and A2 => B5), the fused objects (B1 and B4 fused to a single object and compared to A3) and additionally includes every unmatched cell with a score of 0. In this way, unmatched cells and one-to-one matches are both accounted for and multiplets are included in a comparable manner, with the interface between multiplets (the common boundary between B1 and B4 cells) being neglected since the doublet A3 does not feature this boundary. This makes the complete/modified score the most comprehensive single measure of combined alignment and segmentation quality, since no cell is excluded and multiplets are counted only once. With this score should be stated: the total number of cells, the number of unmatched cells and the number of fused.

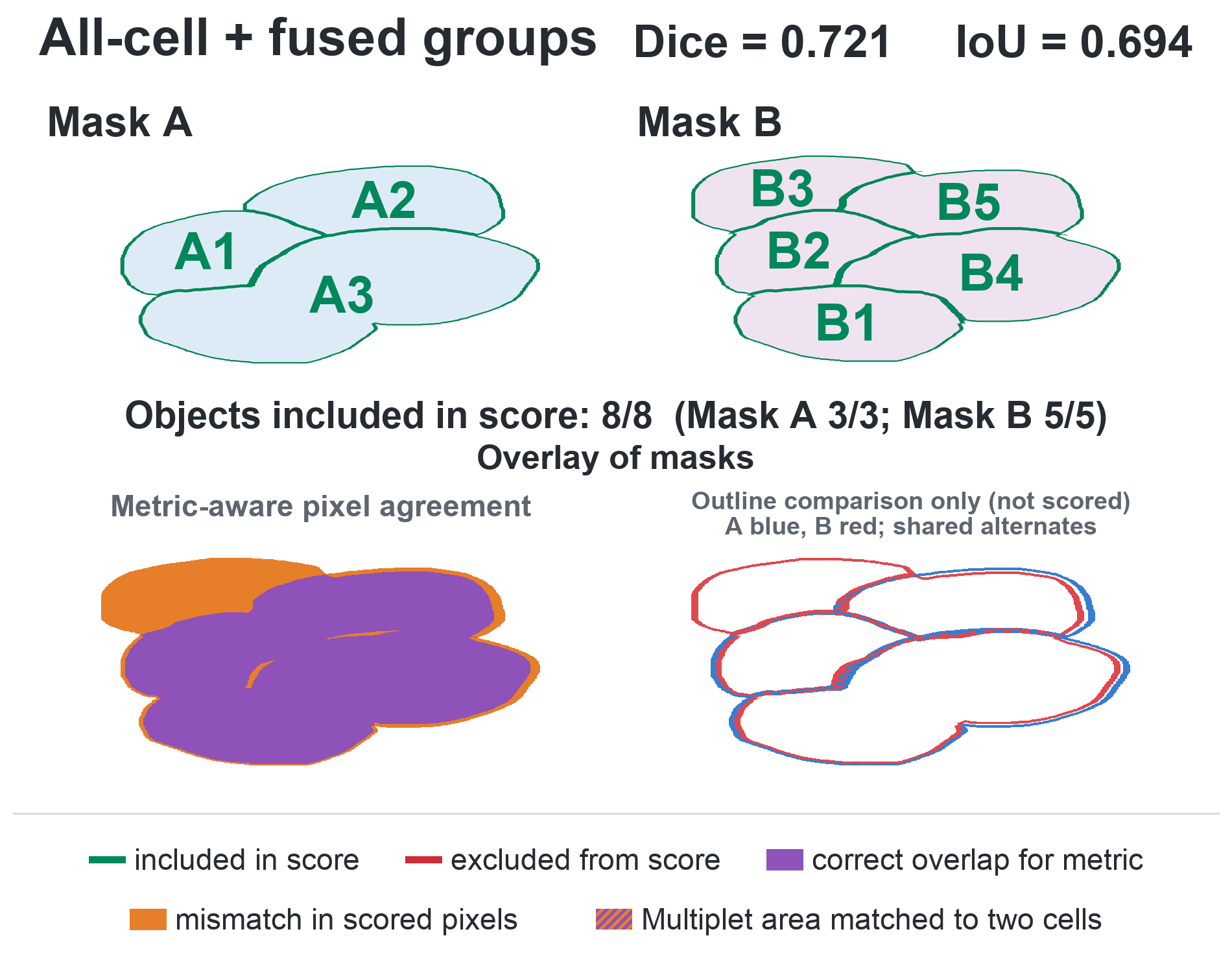

Figure S6: Modified Dice coefficient and IoU.

Figure S*7* illustrates how five Dice/IoU metrics evaluate different aspects of mask agreement. The first four rows show perfectly overlapping masks with identical objects, one missing cell, one additional cell, or a fused-cell group. These examples demonstrate how object-number and segmentation differences affect the scores despite perfect spatial alignment. The final row shows a non-perfect overlap, including differences in outer boundaries and internal cell assignments. Green and red outlines identify objects included in or excluded from each score, while purple, orange, and striped areas indicate correct overlap, mismatch, and multiplet regions matched to two cells, respectively. Together, the metrics provide complementary information about global alignment, cell-level correspondence, and segmentation differences.

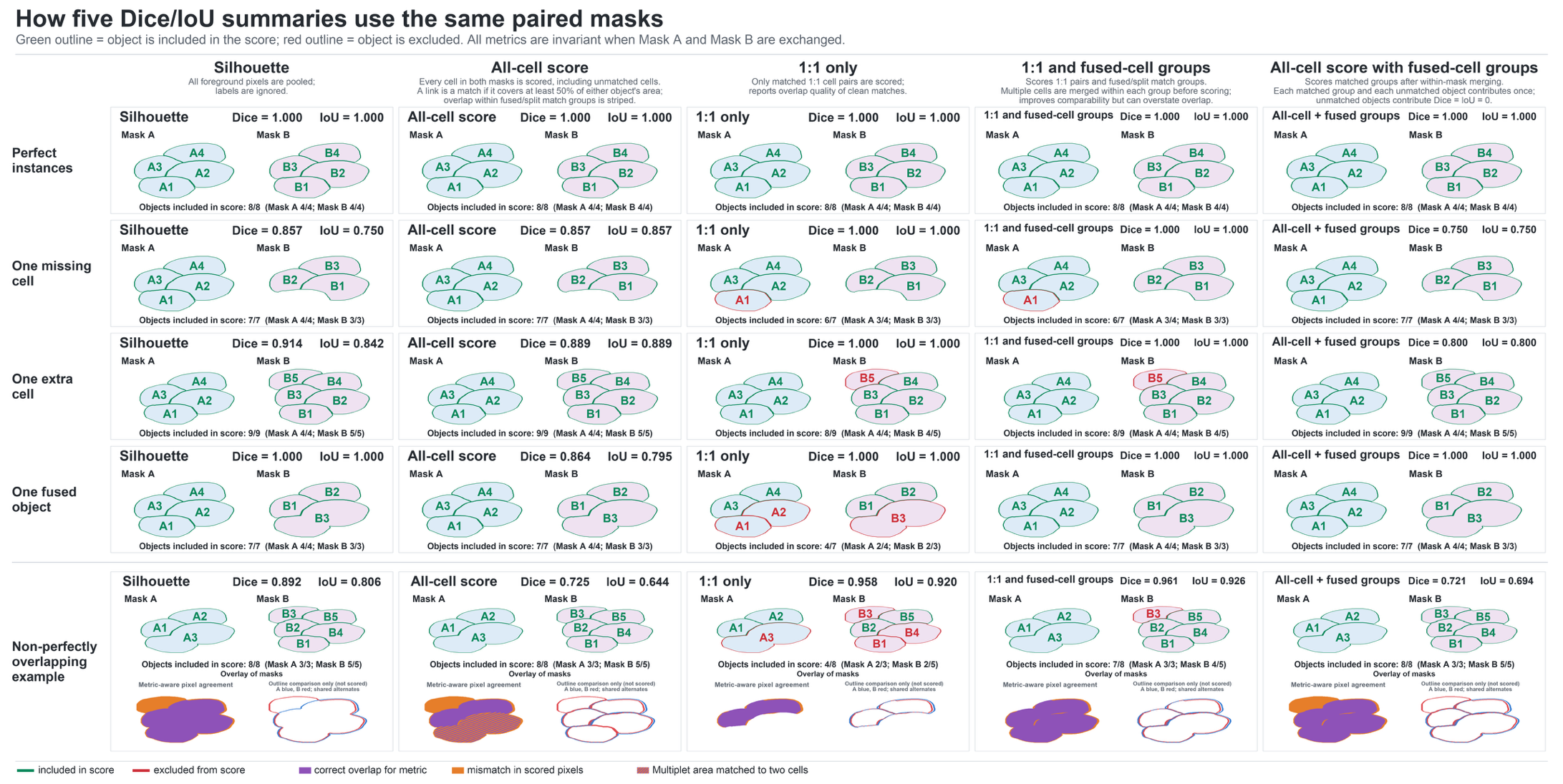

Figure S7: Visual representation of 5 Dice coefficients and IoU scores. Comparison of five Dice and intersection-over-union (IoU) scoring approaches. Columns represent the silhouette, all-cell, 1:1-only, 1:1-and-fused-cell-group, and all-cell-with-fused-cell-group scores. Rows show identical masks, a missing cell, an additional cell, a fused object, and non-perfectly overlapping masks. Green and red outlines indicate objects included in and excluded from the respective score, respectively. In the pixel-level overlays, purple represents correctly matched overlap, orange represents mismatched pixels, and purple–orange stripes indicate a multiplet area matched to two individual cells. In the outline overlays, Mask A is shown in blue and Mask B in red, with coincident outlines displayed as alternating blue and red segments.

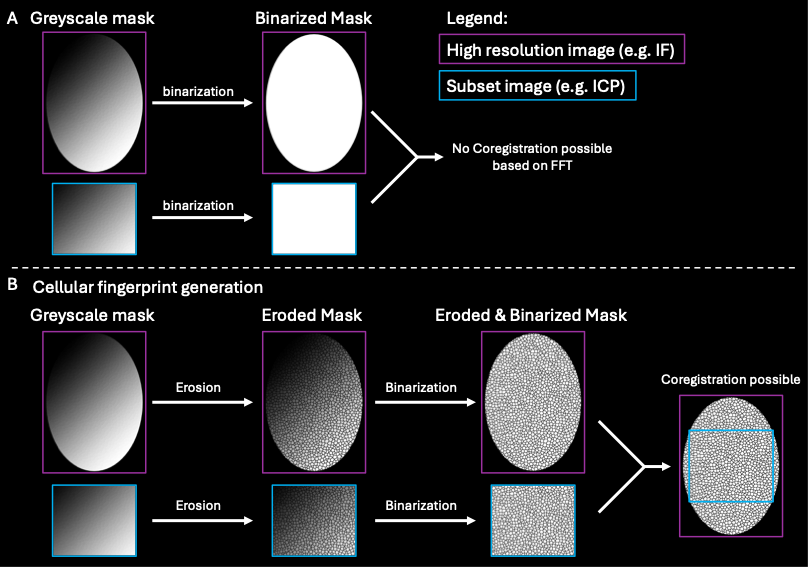

Figure S8: Theoretical example to illustrate the generation of a cellular fingerprint in a sample with no extracellular matrix background (e.g. Tumor). A: Without erosion, registration on a binarized mask without background is not possible. Nor for FFT, nor SDM. B: When firstly eroding each individual cell/object mask, an artificial background is generated, and a cellular fingerprint is generated. This ultimately enables registration based on that fingerprint.

Figure S9: FFT based registration of a Cytospin. The ablated area is a subset of the IF-Image. FFT can locate the correct position. The bounding box of the ICP image plus a padding of 100 pixels in the IF image is cropped (red box) and passed to the gradient mask generation stage. Cropping is performed on the original greyscale mask and not the binarized mask as cellular data is fundamental for stage 2 and 3.

Figure S10: Coregistration of a Cytospin sample containing ten different barcoded experiments. **IF:** FITC-WGA and DAPI were used for segmentation **ICP:** Ir-DNA intercalator and Stacked Anti-WGA signals. **Coregistered Output:** Full and cropped overlay of the FITC-WGA and Stacked Anti-WGA as well as Ir-DNA intercalator and DAPI signals to compare the coregistration. Excellent overlap can be observed as Membranes and DNA match. For visualization purposes, the segmentation masks were colored in a turbo-scale to make cells more visible. Greyscale masks were used for coregistration.

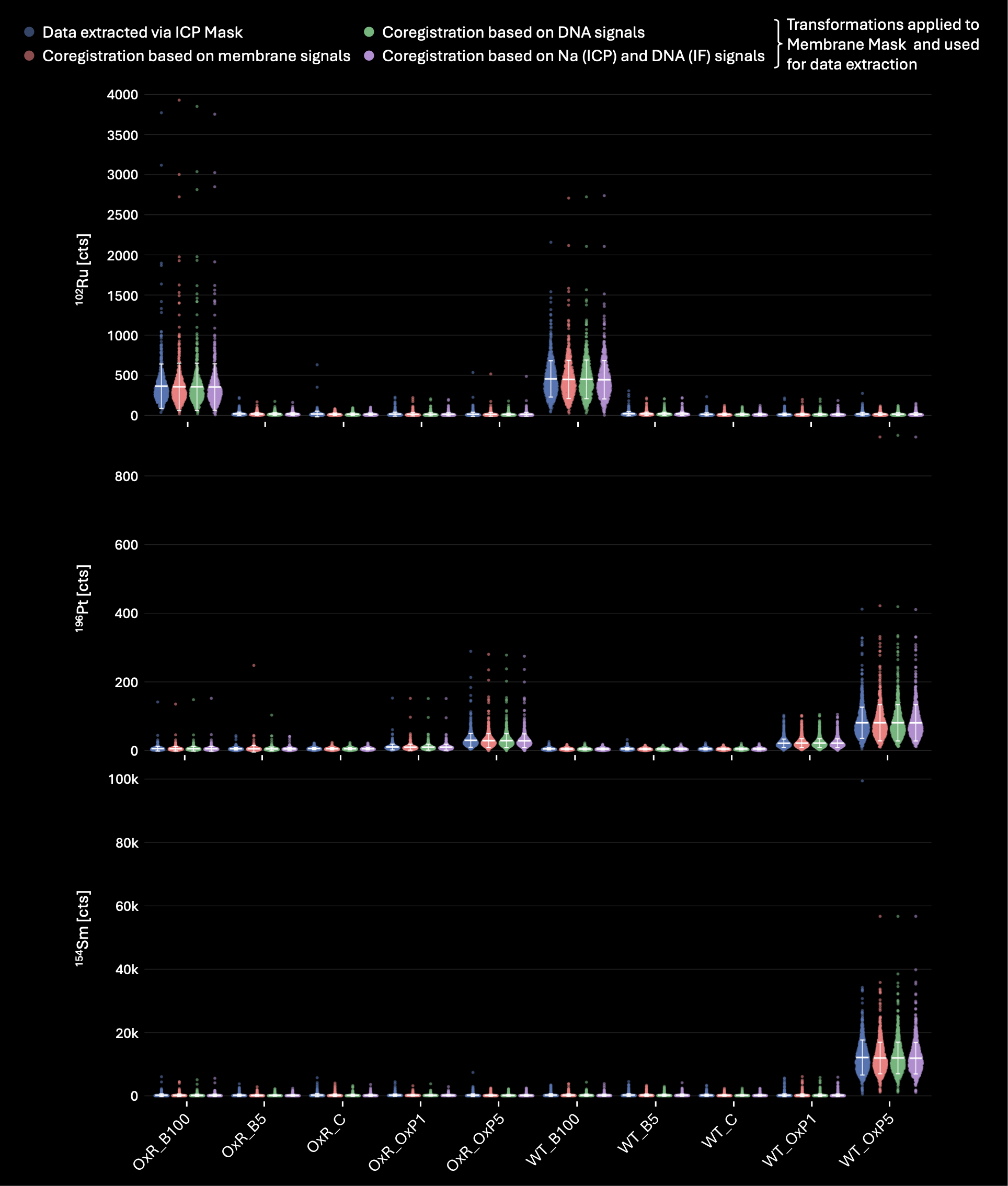

Figure S11: Comparison of Data extracted from the ten barcoded cytospin sample where single cell data was once extracted via ICP membrane-based mask (blue) and three different Coregistration setups. For the red dataset, coregistration was performed based on Membrane based segmentation masks and data extracted via the coregistered IF membrane mask. For the green dataset, coregistration was performed based on the DNA signal-based segmentation masks (IF and ICP). Transformations were applied to the IF membrane mask which was then used for single cell data extraction. For the Violet dataset, coregistration was performed based on a ^23^Na signal-based segmentation masks (ICP) and the DNA signal-based segmentation masks (IF). Transformations were applied to the IF membrane mask which was then used for single cell data extraction. Coregistration was successful with all segmentation masks and means as well as Distributions are in good accordance. Means are listed in Table S9.

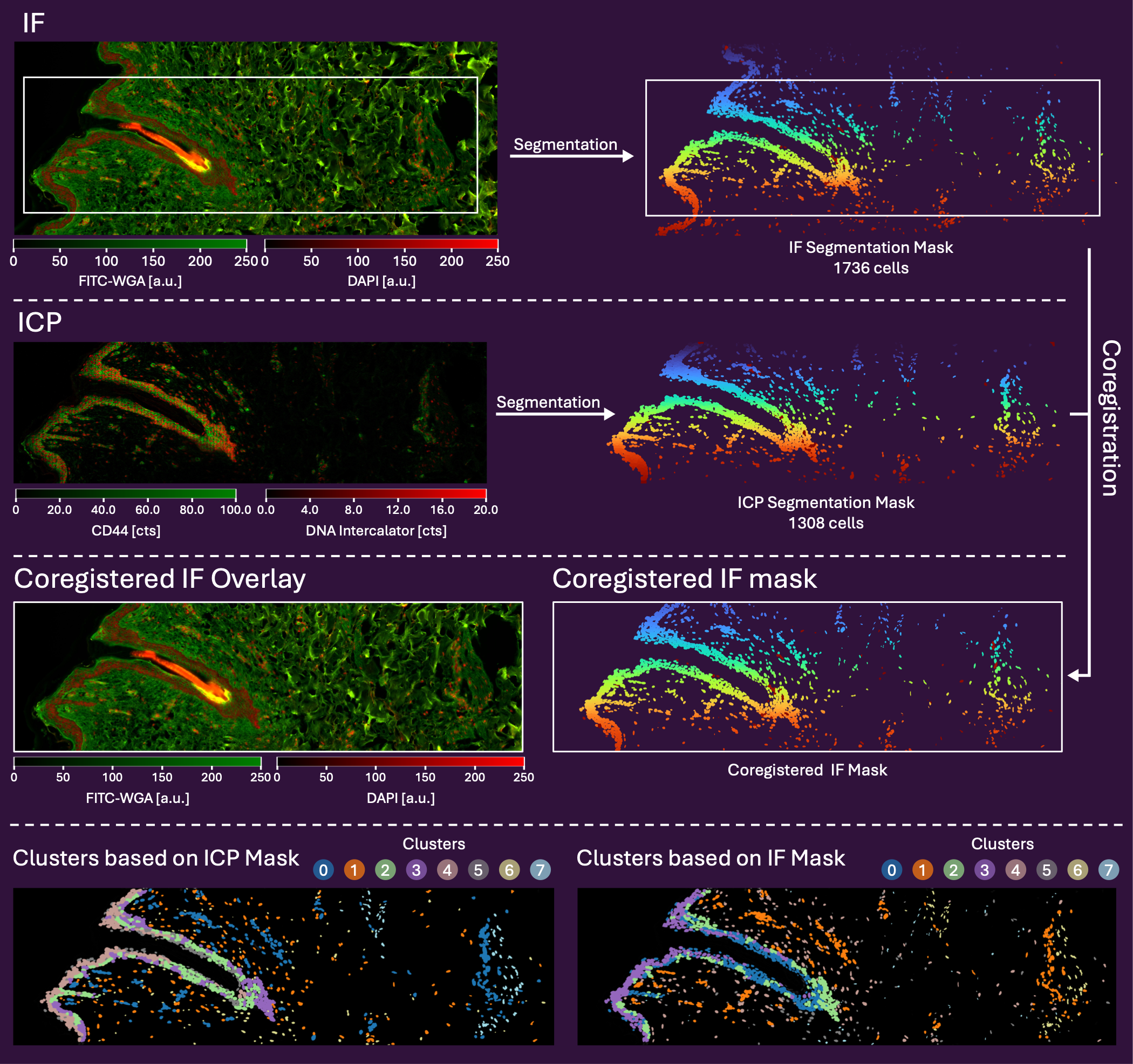

Figure S12: Coregistration between an ex vivo human skin sample analyzed by IF and LA-ICP-TOFMS. Segmentation was performed on the ^193^Ir-DNA-intercalator and CD44 signal for the ICP images and on DAPI and FITC-WGA for IF. White boxes in IF images correspond to the area ablated within LA-ICP-TOFMS. For visualization purposes, the segmentation masks were colored in a turbo-scale to make cells more visible. Greyscale masks were used for coregistration.

**Declaration of generative AI and AI-assisted technologies in the manuscript preparation process**

During the preparation of this work, the authors used ChatGPT (OpenAI, GPT-5 model) and Anthropic (Claude Code, Opus 4.8 and Opus 5) to assist in the development of the coregistration pipeline and for language editing. All scripts generated with AI assistance were independently reviewed, validated, and modified if needed. C.M. critically reviewed and edited the manuscript and takes full responsibility for the content of the published article.
